# False Discovery Rate Control for Grouped Hypotheses: Application to miRNAome Data

**DOI:** 10.1101/2024.01.13.575531

**Authors:** Nilanjana Laha, Salil Koner, Austin Labowitz, Navonil De Sarkar

**Author notes:** Corresponding author: Nilanjana Laha^1^.

## Abstract

Bioinformatics studies often involve numerous simultaneous statistical tests, increasing the risk of false discoveries. To control the false discovery rate (FDR), these studies typically apply a statistical method called the Benjamini–Hochberg (BH) method. However, BH can be overly conservative, particularly in small-sample studies, and it does not take advantage of relevant structural information among the hypotheses, such as groupings. Group structures can arise, for example, when genomic features located in close proximity are co-regulated. Recent statistical developments have yielded group-adaptive BH methods that can leverage pre-existing group information to improve statistical power while maintaining FDR control. However, these methods remain underutilized in bioinformatics practice. In this study, we illustrate the practical application of group-adaptive BH methods using a previously published, moderately scaled microRNA (miRNA) dataset. Even under simple groupings based on chromosomal location, these methods identified more miRNAs with significantly deregulated expression (FDR-adjusted p-value < 0.05) compared to the traditional BH method. Most of the new discoveries are supported by prior literature and a related 2017 study. Although sensitivity to grouping strategy varied across methods, our control analysis indicated that, for most methods, the additional detections may be attributable to the incorporation of group information. Our results highlight the potential of specialized BH methods for controlling the FDR in omics studies with pre-defined group structures, and motivate further evaluation of their generalizability across diverse datasets.

## 1 INTRODUCTION

Molecular profiling technologies, including sequencing, microarrays, and targeted panel assays, enable the simultaneous measurement of many molecular features in biological samples (Dudoit et al., 2003; Goeman and Solari, 2014; Sesia et al., 2021a). Analyzing such data typically involves testing many hypotheses in parallel, each corresponding to one biological element, such as one gene expression in a microarray or RNA-seq study. The rejection of a hypothesis is generally associated with biologically relevant elements, indicating statistical “discovery.” However, testing many hypotheses simultaneously inflates the probability of false positives, making it necessary to apply statistical procedures that control false discoveries (Menyhart et al., 2021). A widely used approach for this purpose is controlling the false discovery rate (FDR) (Korthauer et al., 2019). The FDR is the expected proportion of false rejections among all rejected hypotheses. An FDR of 5% implies that, on average, 5 of 100 significant findings are expected to be false positives.

In the past decades, various statistical methods have been proposed for controlling the FDR. However, the Benjamini and Hochberg (BH) procedure remains the most prominent tool for this purpose in the realms of bioinformatics (Benjamini and Hochberg, 1995; Korthauer et al., 2019). Originally developed for independent hypotheses, the BH method also controls the FDR under weak dependence, an assumption that often holds in biological applications (Benjamini and Yekutieli, 2001). The BH method relies solely on the p-values of the hypothesis tests. It provides a p-value cutoff, and only tests with lower p-values are rejected. Although the BH method reliably controls the FDR, it is known to be conservative when the number of hypotheses greatly exceeds the sample size, which is a common issue in high-dimensional biological studies (Hu et al., 2010; Li and Barber, 2019). The high dimensionality of the data increases the noise within the data, making it harder to distill a weaker signal from the noise. The conservativeness of the BH method can therefore potentially hinder scientific discoveries in such settings.

Additionally, the BH method treats all hypotheses as exchangeable, ignoring any auxiliary biologically information on the hypotheses (Genovese et al., 2006). In many domains, including bioinformatics, such information can be available. This study focuses on hypotheses with pre-existing group structures, which often arise naturally in biological data. For example, closely located genes are frequently correlated and may be co-expressed due to shared transcription programs and regulatory mechanisms, forming positional clusters (Koutná et al., 2007).

Recent advancements in statistics have led to modified BH methods that can leverage pre-existing group structures, e.g., the grouped BH (GBH) methods of Hu et al. (2010) and the structure adaptive BH algorithm (SABHA)^1^ of Li and Barber (2019). Such methods recalibrate the p-values, prioritizing rejections from potential critical groups. A group’s criticality is determined by the proportion of true null hypotheses within it; smaller proportions indicating greater criticality. When pre-existing group structure is present, these *group-adaptive BH* methods were shown to achieve higher statistical power than the BH method in prior controlled experiments (Korthauer et al., 2019; Hu et al., 2010; Li and Barber, 2019). A higher statistical power indicates the ability to detect lower signals, thus potentially increasing the number of scientific discoveries. Nevertheless, these group-adaptive BH methods have yet to see widespread adoption in bioinformatics practice.

Many other FDR control methods are available for false discovery rate control with potentially dependent hypotheses, e.g., independent hypothesis weighting (Ignatiadis et al., 2016), q-values (Storey, 2002), etc. An emerging statistical technique called knock-off has also shown promise for FDR control and is flexible enough to incorporate group structures (Sesia et al., 2021a). However, most of these methods are typically designed to handle broad classes of dependence structures and are not specific to pre-existing group structures (Li and Barber, 2019; Friston, 2003). Moreover, methods such as independent hypothesis weighting require covariates independent of the p-values (Ignatiadis et al., 2016). Since our primary aim is to illustrate how grouping, motivated by domain knowledge, improves power while controlling the FDR, we restrict our attention to group-adaptive BH methods, especially GBH and SABHA, to keep the scope of the study coherent and allow for a clearer interpretation of the results. Notably, these methods do not use groups as the unit of rejection. Instead, rejections are conducted at the level of the individual hypotheses. However, FDR control methods that reject hypotheses at the group level are also available (Efron and Tibshirani, 2002; Benjamini and Heller, 2007).

This work is an applied study illustrating the integration of group-adaptive BH procedures into moderate-throughput microarray analyses with pre-specified group structures. To this end, we use the microRNA (miRNA) dataset from De Sarkar et al. (2014), in which one of the present co-authors was involved. The dataset comprises 522 miRNAs measured using a targeted qPCR panel from 18 cancer patients with gingival buccal squamous cell carcinoma (GBSCC), a subtype of oral squamous cell carcinoma (OSCC). Although modest in scale compared to modern RNA-seq experiments, this dataset is still high-dimensional, with the number of assays far exceeding the sample size, and is representative of settings where targeted panels are applied to limited cohorts. Addressing the challenge of small sample size (*n* = 18) required careful analysis choices, and the pipeline presented here may serve as a useful template for similar studies. Such contexts remain common in translational and clinical studies, making them valuable for illustrating the practical utility of group-adaptive BH methods (Bustin et al., 2009). Because the discussed procedures require only pre-specified group structures, the encouraging results from this dataset suggest that they may also be useful in larger-scale, high-dimensional bioinformatics studies where similar group information is available.

De Sarkar et al. (2014)’s goal was to investigate the role of miRNAs in GBSCC by comparing their expression in histopathologically confirmed malignant and healthy normal (control) tissues. As there were 522 miRNA assays, 522 t-tests were performed. The BH method was used to control the FDR at 5%, and seven miRNAs whose expression were significantly deregulated in the malignant tissues were identified. However, De Sarkar et al. (2014) noticed that the expression levels of some cancer biomarkers, including hsa-miR-21 and hsa-miR-1, were either borderline or below the BH threshold. Some of the target genes of these miRNAs were found to be significantly deregulated by Singh et al. (2017) in a follow-up whole transcriptome analysis study, which analyzed a case series substantially overlapping with that of De Sarkar et al. (2014). This observation raised the question of whether the BH method had sufficient statistical power to detect deregulated miRNAs in their dataset. De Sarkar et al. (2014)’s dataset therefore serves as a suitable case study for evaluating whether group-adaptive FDR methods can improve the detection of biologically relevant miRNAs.

Considering our limited sample size (*n* = 18), the number of miRNAs analyzed (*p* = 522), and their distribution across the genome, we opted for a relatively straightforward grouping approach in this proof-of-concept analysis. Because proximal miRNAs have been reported to exhibit co-expression (Koutná et al., 2007), the physical location of the miRNAs on the chromosome was used as the basis for grouping. While overly simplistic, this strategy was suitable for the small sample size and serves our purpose of illustrating how incorporating external biological information can improve detection. Even under this simple criterion, multiple grouping schemes are possible. We compared several such schemes, defined solely by genomic position and independent of expression data. Then we demonstrated the application of group-adaptive BH methods under these schemes and compared their performance with the BH method. A small control study was also conducted using randomly assigned groups. Existing literature and the whole transcriptome analysis by Singh et al. (2017) are used to discuss the relevance of the additional miRNAs discovered by the group-adaptive BH methods.

## 2 MATERIAL AND METHODS

Section 2.1 provides a brief overview of the study population and the resulting miRNAome dataset. Sections 2.2 and 2.3 describe the implementation of the *t*-tests and the group-adaptive BH methods, respectively. Section 2.4 discusses the various position-based schemes considered for grouping the miRNAs.

### 2.1 Study participants and miRNAome dataset

We used publicly available data from De Sarkar et al. (2014). The study participants were 18 unrelated Indians aged 39-80 years with tobacco habits, with a male:female ratio of 5: 4. All patients had histopathologically confirmed GBSCC, which is a type of OSCC prevalent in the tobacco-chewing population in South Asia. The dataset contained ΔCt values for 522 miRNAs derived from 18 tumor-normal sample pairs of the 18 patients. ΔCt values serve as surrogate estimates for relative miRNA expression levels (relative to geometric mean expression of three endogenous control genes). The ΔCt values were processed to obtain the ΔΔCt values, where ΔΔCt = ΔCt of a miRNA in cancer tissue ™ ΔCt of that miRNA in control tissue. Further details on the data collection methods are provided in Section S1 of the Supplement.

#### 2.1.1 Missingness

Some miRNAs were not expressed in all 18 patients in De Sarkar et al. (2014)’s dataset. In their study, miRNA expression profiling was performed using TaqMan Low-Density Arrays (TLDA-A V2 and TLDA-B V3) on the 7900HT FAST Real-Time PCR System (Applied Biosystems, USA), equipped with a TLDA flat block. This platform is adequately sensitive, but similar to all qPCR-based methods, it has a fixed lower detection limit. If either the tumor or the matched normal expression value for a given miRNA fell below this detection threshold, De Sarkar et al. (2014) excluded both values to simplify downstream analyses. Consequently, some miRNA pairs were missing in the miRNA dataset. While this lower detection limit is likely a contributor to the missingness, other technical or biological factors may also have played a role. Nonetheless, the missingness in the miRNAome measurements appears to be not at random (MNAR) (Hughes et al., 2019).

Each patient had at least 12 missing miRNA expression pairs, although most had fewer than 50 missing pairs. Supplementary Figure S2.1 displays the number of missing miRNA pairs for each patient, and Supplementary Figure S2.2 shows the histogram of the number of missing pairs across patients. Overall, 59.77% of miRNAs had at least one missing tumor–normal pair, and 30% had more than two missing pairs. For further details on missingness, refer to Section S2 in the Supplement.

De Sarkar et al. (2014) imputed the missing observations using an imputation strategy based on sample medians in their statistical analysis. However, we observed a sharp increase in the number of discoveries regardless of the FDR control method when using simplistic imputation strategies such as median or mean imputation. This is unsurprising because it is well-known that imputation with the median or mean can artificially decrease the sample variance, especially when the sample size is small, leading to smaller p-values (Zhang, 2016; Sarkar and Zhao, 2022). This can increase the risk of false discoveries for test units both with or without missing values. Both the BH procedure and its variants rely on ranking p-values to determine an adjusted p-value threshold. When imputation results in smaller p-values, the adjustment mechanism can yield a larger overall threshold (cf., e.g., Benjamini and Hochberg, 1995, for details on the adjustment). For example, in our dataset, the BH-corrected p-value threshold increased from 0.00020 in the non-imputed data to 0.00094 under median imputation. This means that, in the non-imputed data, BH considered miRNAs with p-value ≤ 0.00020 as significantly deregulated, whereas after imputation, BH considered any miRNA with p-values ≤ 0.00094 to be significantly deregulated.

A more sophisticated imputation strategy may reduce the risk of imputation-driven false detections. However, given our limited sample size, the relatively high proportion of missing values, and the complexity of measuring all biological factors behind miRNA expression, we chose not to impute missing values. Instead, we conducted our statistical analysis using only complete cases. Hughes et al. (2019) showed that complete case analysis can be statistically valid in MNAR data. Finally, since our groups are informed by external biological information, they are unaffected by data missingness.

### 2.2 Pairwise t-tests

The t-tests were conducted as in De Sarkar et al. (2014); details are provided here for completeness. Up-regulation or downregulation of miRNA expression was determined by checking if the change of miRNA expression exceeded 2 fold compared to the control. The fold-changes are measured by ΔΔCt value: a positive ΔΔCt implies *>*2 fold downregulation, while a negative ΔΔCt implies *>*2 fold upregulation of miRNA expression. We performed 522 one-sided pairwise t-tests to determine whether the expression of miRNAs was significantly deregulated. For each miRNA, the direction of the one-sided paired t-test was determined by the sign of the median ΔΔCt value across the 18 patients. The null hypothesis of the one-tailed paired t-test was that the expression of a particular miRNA is not greater than 2 fold upregulated (or downregulated).

### 2.3 Group-adaptive BH methods

As noted in Section 1, group-adaptive BH methods assign greater importance to groups with a higher estimated proportion of true null hypotheses. The group-wise null proportions can be estimated using various statistical approaches. Examples include the two-stage step-up (TST) method (Benjamini et al., 2006), the Least-Slope (LSL) method (Benjamini and Hochberg, 2000), and likelihood-based approach (Li and Barber, 2019), which lead to TST-GBH, LSL-GBH, and the group-aware version of SABHA, respectively (Hu et al., 2010; Li and Barber, 2019). As mentioned in Section 1, we focus on these three group-adaptive BH methods and compare them with the standard BH procedure. When the p-values are independent or weakly correlated, these methods were shown to control the FDR if group-wise null proportions are reasonably well-estimated, (Hu et al., 2010; Li and Barber, 2019). TST-GBH and LSL-GBH methods were previously applied by Zhang et al. (2013) to discover differentially methylated regions in the human genome and by Liu et al. (2015) to discover differentially expressed unigenes during the blooming process of Asteraceae flowers.

#### Implementation

TST-GBH and LSL-GBH were implemented following the algorithms provided by Hu et al. (2010). To implement SABHA, we used the R codes provided by the authors of the corresponding paper by Li and Barber (2019). SABHA requires two tuning parameters: (1) *ε*, a lower bound on the proportion of the true null hypotheses, and (2) a threshold *τ*, used for calibrating the individual p-value thresholds. Similar to Li and Barber (2019), we used *ε* = 0.1 and *τ* = 0.5. We found that the discoveries were not sensitive to small perturbations of either of these tuning parameters. We implemented BH using the package Stats in the software R. Our code is provided at the GitHub repository Koner et al. (2023).

### 2.4 Grouping schemes

As noted earlier, miRNAs were grouped by chromosomal location. While co-expression may arise from other biological factors, this simple scheme was appropriate for the small sample size (*n* = 18) relative to the number of genes (∼ 500) in this proof-of-concept analysis. The miRNAs can also be grouped using data-dependent clustering methods such as k-means clustering, tight clustering (Karmakar et al., 2019), and weighted gene co-expression network analysis (WGCNA) (Horvath, 2011). However, recent studies highlighted the risk of potentially elevated type-I error when using the same data for clustering and downstream testing (Gao et al., 2020a; Luecken and Theis, 2019; Lähnemann et al., 2020). Given our limited sample size, reusing the miRNA expression data for group assignment would further exacerbate this risk.

Even when grouping miRNAs by spatial location on the chromosome, multiple grouping schemes/ strategies are possible. We considered hierarchical schemes, starting with the coarsest scheme, in which miRNAs were grouped solely by chromosome number, and progressively splitting the groups to obtain finer schemes. The first three schemes were: (a) grouping by chromosome number (miRNAs on the same chromosome form one group); (b) grouping by chromosome number and strand (miRNAs on the same chromosome and strand form one group); and (c) grouping by chromosome number, strand, and arm (miRNAs on the same chromosome, strand, and chromosomal arm form one group). Groups with no miRNAs among the 522 assayed were excluded, and single-miRNA groups were merged with the nearest neighboring group.

To obtain finer groupings from Scheme (c), we generated multiple new grouping schemes by splitting large groups in Scheme (c) according to different maximum group sizes, where group size refers to the number of miRNAs in a group. Specifically, for each *k* ∈ {25, 23,…, 5}, we formed a new scheme in which any Scheme (c) group with more than *k* members was divided into adjacent subgroups of size *k*. The chromosome coordinates provided by De Sarkar et al. (2014) were used to guide splitting, ensuring that adjacent miRNAs were grouped together. If the group size was not a multiple of *k*, the final subgroup contained fewer than *k* members. For simplicity, we may refer to this family of refinements as “grouping schemes derived from Scheme (c)”.

As the grouping schemes becomes progressively finer from Scheme (a) to *k* = 5, the total number of groups per scheme increases accordingly. Figure 1 plots the total number of groups for each scheme. Schemes (a), (b), and (c) have 23, 44, and 67 groups, respectively, rising to 129 at *k* = 5. We did not consider schemes with *k* ≤ 4 because the number of groups becomes excessively large. For instance, *k* = 2 yields 278 groups. This is problematic because, as we shall see in Section 3, the GBH methods tend to become too liberal when the number of groups is large.

**Figure 1.**
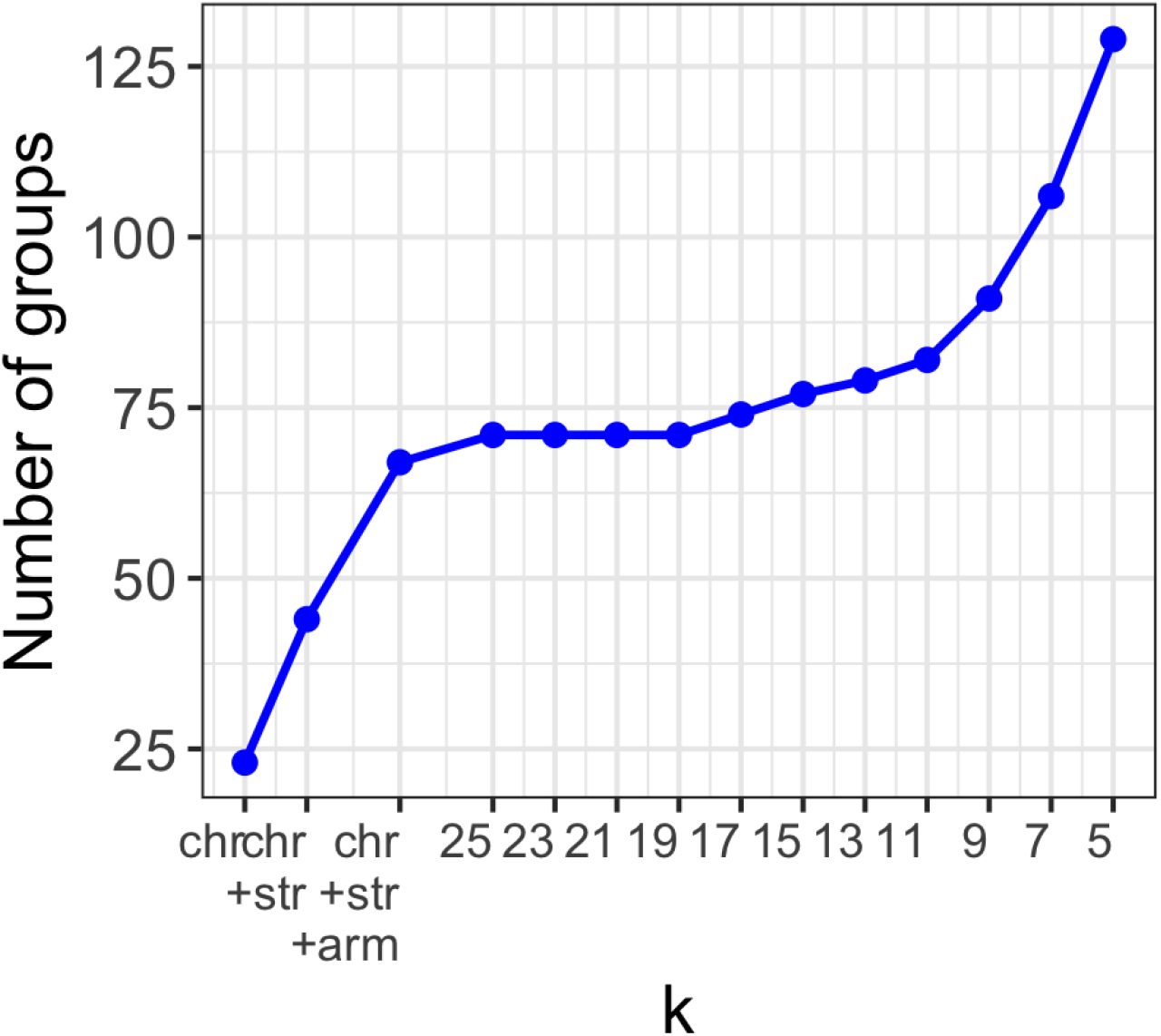
Plot of the total number of groups as a function of grouping schemes. The grouping schemes were as follows: Chr refers to Scheme (a) or grouping by chromosome number only; Chr+Str refers to Scheme (b) or grouping by chromosome number and strand; and Chr+Str+Arm refers to Scheme (c) or grouping by chromosome number, strand, and arm. For each *k*, any group from Scheme (c) with more than *k* miRNAs was split into adjacent groups of size *k*. If the group size was not a multiple of *k*, the final group had fewer than *k* miRNAs. The miRNAs’ coordinates on the chromosome, provided by De Sarkar et al. (2014), were used to guide this splitting so that adjacent miRNAs were grouped together. The Y-axis represents the total number of groups resulting from each grouping scheme.

Table S3.1 in the Supplement tabulates the sizes and locations of the groups in Scheme (c). Figure S3.1 in the Supplement displays the histograms of the group sizes for Schemes (a), (b), (c), and when *k* = 15 and 5. Figure 1 indicates that the number of groups changes little when Scheme (c) is split into finer groups up to *k* = 11, but increases sharply when *k* falls below 11. This occurs because Scheme (c) contains only a few large groups. Eleven groups, which is about only 16% of all groups in Scheme (c), have size ≥ 11. In contrast, this scheme contains a larger proportion of moderately sized groups, with 17 groups (about 25%) with size ≥ 9 (see also Table S3.1 and Figure S3.1iii).

#### 2.4.1 Intra- and inter-group association

Hughes et al. (2019) observed that group-adaptive BH methods exhibit higher statistical power than the standard BH method when the intra-group association, that is, the association among test units within the same group, is stronger than the inter-group association, which refers to the association between test units in different groups. Visual inspection of pairwise correlations between miRNAs’ ΔΔCt values suggests that correlations within groups are generally higher or comparable to those across groups. Figure S3.2 in the Supplement illustrates examples with two Scheme (c) groups.

For each of our grouping schemes, we heuristically compared the intra-group and inter-group correlations between the ΔΔCt values utilizing a simple random effect model, which is a widely used statistical approach (Nakagawa et al., 2017). Details on fitting this model and the corresponding quantification of inter-group and intra-group correlations are provided in Appendix 5. The simple random effect model assumes the inter-group and intra-group correlations to be constant across all groups. Even if this assumption is violated, the estimated correlations provide meaningful assessments of the overall strength of the inter-group and intra-group association. Figure S3.3 in the Supplement shows the estimated correlations across our grouping schemes. The estimated inter-group correlation ranged between 0.041 and 0.044 for all grouping schemes, indicating a weak inter-group association. In contrast, the intra-group correlation estimates were 0.108, 0.119, and 0.139 for schemes (a), (b), and (c), respectively, and ranged between 0.147 and 0.18 for all other grouping schemes considered in this study. Thus, the simple random effect model suggested a weaker inter-group association compared to intra-group association for our grouping schemes. Figure S3.3 in the Supplement also shows that the intra-group correlation estimates generally increase with finer grouping schemes. This is expected because intra-group correlation tends to increase when within-group variation decreases, which occurs as the grouping becomes finer (Searle et al., 2009).

## 3 RESULTS

Section 3.1 presents detection patterns of the group-adaptive BH methods under different grouping schemes. Section 3.2 reports a control study with randomly assigned, biologically uninformative groups. Section 3.3 discusses the relevance of the newly identified miRNAs in cancer.

### 3.1 Significantly deregulated miRNAs

In this section, we compare the significantly deregulated miRNAs detected by BH, TST-GBH, LSL-GBH, and SABHA. All methods were applied at 5% level of significance. BH identified three miRNAs, hsa-miR-133a-3p, hsa-miR-31-3p, and miR-206, as significantly deregulated. The detections of the group-adaptive BH methods depended on the grouping scheme but always included these three miRNAs. Figure 2 shows the number of significantly deregulated miRNAs for each method across grouping schemes. TST-GBH was the most liberal of the four methods, yielding the highest number of detections for any scheme, consistent with the observations of Sarkar and Zhao (2022). The number of detections by TST-GBH and SABHA generally increased as the grouping scheme became finer, whereas LSL-GBH showed no consistent trend.

**Figure 2.**
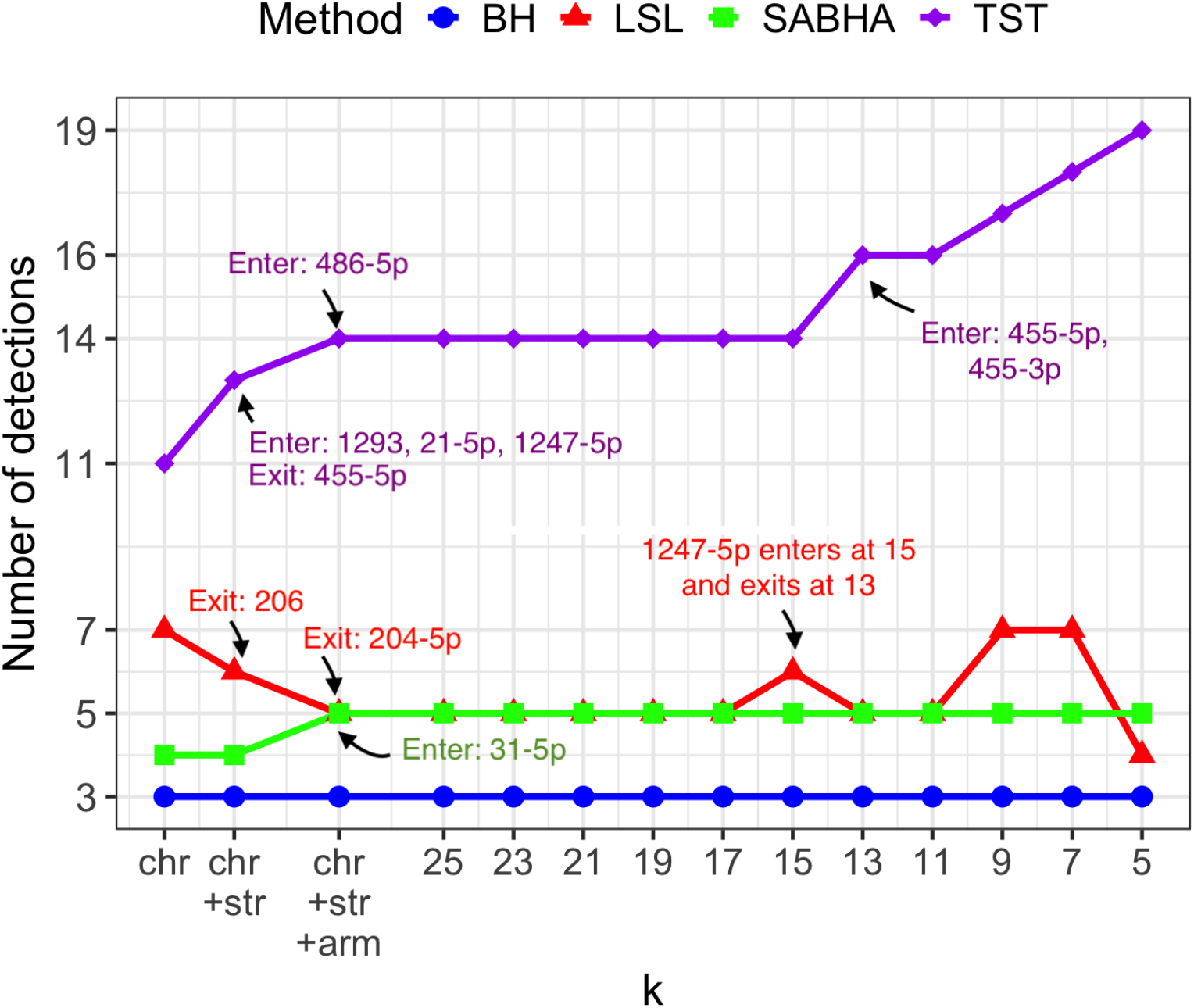
Plot of the number of detections by each FDR control method vs grouping scheme. The methods compared were BH, TST-GBH (abbreviated as TST), LSL-GBH (abbreviated as LSL), and SABHA, each applied with an FDR control level of *α* = 5%. The first three grouping schemes were: (a) grouping by chromosome number only (Chr); (b) grouping by chromosome number and strand (Chr+strand); and (c) grouping by chromosome number, strand, and arm (Chr+Str+Arm). For each *k*, any group from Scheme (c) with more than *k* miRNAs was split into adjacent groups of size *k*, using the miRNAs’ chromosome coordinates from De Sarkar et al. (2014) to ensure adjacent miRNAs were grouped together. If the group size was not a multiple of *k*, the final group contained fewer than *k* miRNAs.The Y-axis shows the total number of detections from each grouping scheme. Detection sets changed little from one scheme to the next, except for a few miRNAs entering or leaving. When a new miRNA entered a detection set, its name is listed under “Enter”; when an miRNA exited, its name is listed under “Exit.” Since BH is not group-dependent, there are no entries or exits. A full list of miRNAs detected as significantly deregulated under Scheme (c) and finer schemes appears in Table 1.

Among the group-adaptive BH methods, SABHA was the most conservative and most robust to changes in grouping schemes, followed by LSL-GBH. As shown in Figure 2, SABHA’s detections were the most similar to BH’s among the three group-adaptive methods. Under the two coarsest schemes, i.e., Schemes (a) and (b), SABHA’s detections differed from that of BH only by the inclusion of hsa-miR-1. Under Scheme (c) and the finer schemes derived from it, only hsa-miR-31-5p joined SABHA’s detections. Figure 2 shows that the set of significantly deregulated miRNAs varied minimally for all methods across the derived grouping schemes when the maximum group size *k* was in {11, 13,…, 25}, differing by at most two miRNAs. This stability is unsurprising because the group compositions are nearly identical for *k* ∈ {11, 13,…, 25} (see Section 2.4). In contrast, when the maximum group size dropped below 10, detections by TST-GBH—and, to a lesser extent, LSL-GBH—became more sensitive to the choice of grouping scheme. This probably occurs because, in this range, both the number of groups and the estimated intra-group correlation rise more sharply with decreasing *k* (Figure 2; supplementary Figure S3.3).

**Table 1.** Table of significantly deregulated miRNAs under grouping Scheme (c) and the derivative schemes. The FDR control methods are BH, TST-GBH (TST), LSL-GBH (LSL), and SABHA. The listed miRNAs were detected as significantly deregulated (5% significance level) by at least one method when the grouping scheme was either Scheme (c) or a derivative scheme obtained from it. In Scheme (c), miRNAs were grouped by chromosome number, strand, and arm. The derivative schemes considered here have maximum group sizes *k* ∈ {25, 23,…, 11}. In a derivative scheme for a given *k*, any Scheme (c) group with more than *k* miRNAs was split into adjacent subgroups of size *k*, using chromosome coordinates from De Sarkar et al. (2014) to ensure that adjacent miRNAs remained together. If the group size was not a multiple of *k*, the final subgroup contained fewer than *k* miRNAs. If a method detected a miRNA under all grouping schemes, its row is marked with “x”; otherwise, the row lists the schemes where it was detected. The “Chr.” column indicates the chromosome number; the “strand” column indicates the transcribing strand; and the “arm” column indicates the arm of the miRNA gene. The ΔΔCt values are used as surrogate estimates of differential expression for each miRNA. As TST-GBH, LSL-GBH, and SABHA do not yield FDR-corrected p-values, we have reported the raw p-values before FDR correction. The rows are sorted according to p-value. The arrow signs after the p-values indicate whether the expression of the miRNAs was upregulated (↑) or downregulated (↓) (per the sign of ΔΔCt). The “Missing pairs” column shows the number of missing observations for each miRNA. The expressions of miRNAs superscripted with an asterisk were reported to be deregulated by De Sarkar et al. (2014).

| miRNA | Chr. | Str. | Arm | $\Delta\Delta Ct$ | p-value | Miss<br>-ing<br>pairs | BH | TST | LSL | SABHA |
| --- | --- | --- | --- | --- | --- | --- | --- | --- | --- | --- |
| hsa-miR-133a-3p* | 18 | - | q | 6.7 | $5.32E^{-05} \downarrow$ | 0 | x | x | x | x |
| hsa-miR-31-3p* | 9 | - | p | -3.8 | $1.31E^{-04} \uparrow$ | 2 | x | x | x | x |
| hsa-miR-206* | 6 | + | p | 6.0 | $1.97E^{-04} \downarrow$ | 0 | x | x | | x |
| hsa-miR-31-5p* | 9 | - | q | -3.4 | $6.18E^{-04} \uparrow$ | 0 | | x | x | x |
| hsa-miR-204-5p* | 9 | - | q | 4.6 | $8.81E^{-04} \downarrow$ | 2 | | x | | |
| hsa-miR-1 | 18 | - | q | 5.2 | $9.38E^{-04} \downarrow$ | 0 | | x | x | x |
| hsa-miR-7-5p* | 15 | + | q | -3.1 | $9.50E^{-04} \uparrow$ | 2 | | x | x | |
| hsa-miR-1293* | 12 | - | q | -4.8 | $2.77E^{-03} \uparrow$ | 9 | | x | | |
| hsa-miR-486-3p | 8 | - | p | 2.4 | $2.81E^{-03} \downarrow$ | 0 | | x | | |
| hsa-miR-21-5p | 17 | + | q | -2.2 | $3.67E^{-03} \uparrow$ | 0 | | x | | |
| hsa-miR-147b | 15 | + | q | -2.1 | $5.52E^{-03} \uparrow$ | 2 | | x | | |
| hsa-miR-99a-3p | 21 | + | q | 2.8 | $5.83E^{-03} \downarrow$ | 0 | | x | | |
| hsa-miR-455-3p | 9 | + | q | -1.7 | $5.98E^{-03} \uparrow$ | 0 | | $k = 11, 13$ | | |
| hsa-miR-1247-5p | 14 | - | q | 2.4 | $8.93E^{-03} \downarrow$ | 2 | | x | $k = 15$ | |
| hsa-miR-455-5p | 9 | + | q | -1.4 | $1.30E^{-02} \uparrow$ | 0 | | $k = 11, 13$ | | |
| hsa-miR-486-5p | 8 | - | p | 2.0 | $1.78E^{-02} \downarrow$ | 0 | | x | | |

In view of the above, we restrict our focus to miRNAs detected as significantly deregulated by at least one FDR control method under schemes (a), (b), (c), or any derived scheme with *k* ∈ {11, 13,…, 25}. This yields a total of sixteen miRNAs, listed in Table 1. Among these significantly deregulated miRNAs, hsa-miR-31-3p/5p, miR-7-5p, miR-21-5p, miR-147b, miR-1293, and miR-455-3p/5p were upregulated, while miR-133a-3p, miR-206, miR-204-5p, miR-1, miR-486-3p/5p, miR-99a-3p, and miR-1247-5p were downregulated. The list of miRNAs detected under Scheme (a) or Scheme (b) is provided in Table S4.1 in the Supplement. Figure 3 provides a heatmap of the ΔΔCt values of these significantly deregulated miRNAs.

**Figure 3.**
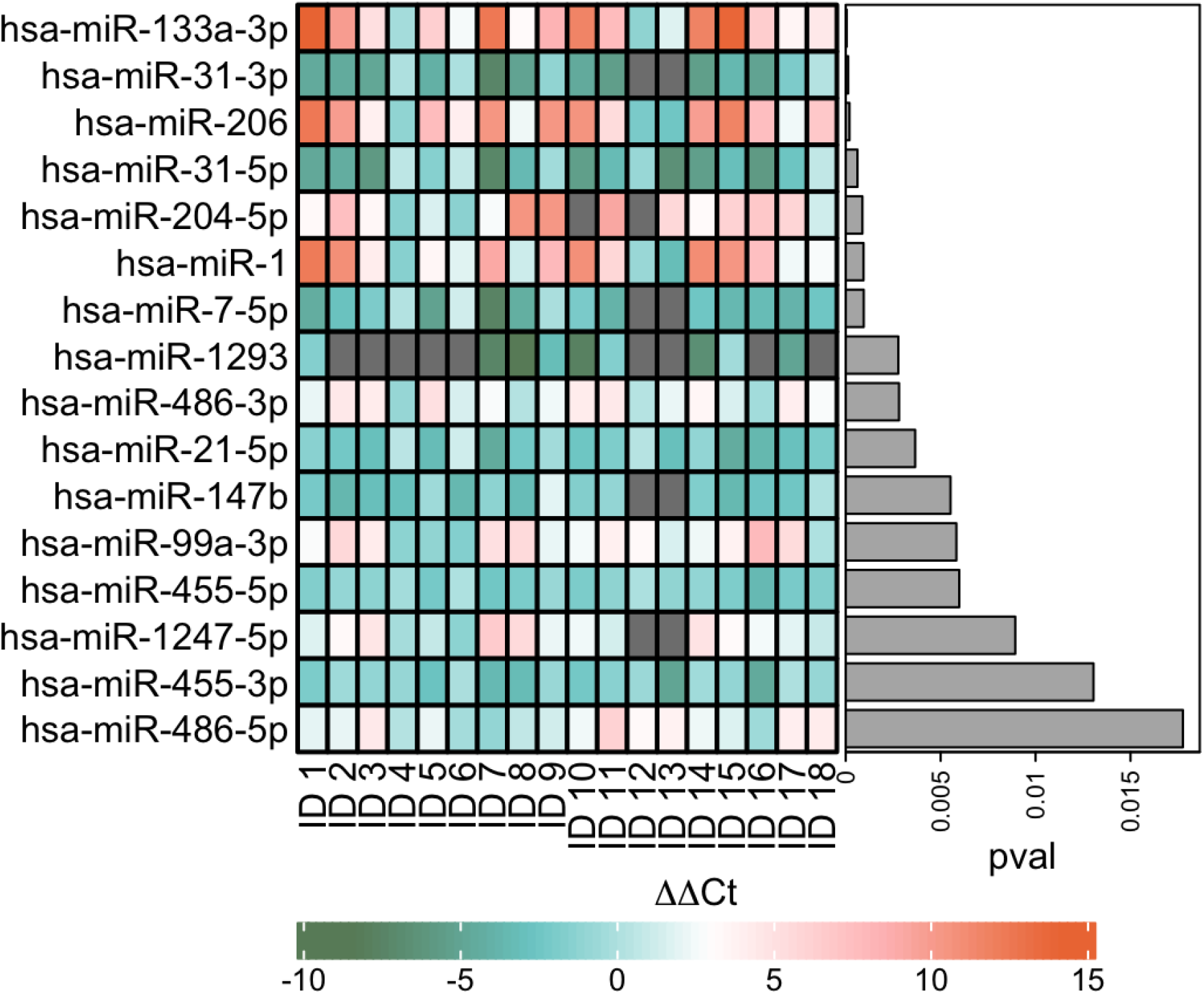
Heatmap of the ΔΔ*Ct* values of the deregulated miRNAs in in Table 1. The heatmap displays deregulation patterns of miRNA expression (corresponding to the miRNAs in Table 1) in the cancer tissues derived from 18 GBSCC patients. The miRNAs are ordered as per unadjusted p-values (indicated as “pval”) obtained from the paired t-tests.

#### Comparison with De Sarkar et al. (2014)

Amomg the sixteen miRNAs listed in Table 1, hsa-miR-133-3p, miR-206, miR-31-3p/5p, miR-1293, miR-7-5p, and miR-204-5p were identified as significantly deregulated in De Sarkar et al. (2014)’s analysis. De Sarkar et al.’s bioinformatics analyses showed that, except for miR-1293, the deregulation patterns of these miRNAs were concordant with previously published studies in OSCC. Although De Sarkar et al. (2014) used the BH method to control the FDR, they identified seven miRNAs instead of three because they imputed the missing values using a median-based imputation strategy. As discussed in Section 2.1.1, such imputations can increase detections by artificially deflating the sample variance. Figure S4.1 in the Supplement shows that the number of discoveries substantially increases for all four FDR-control methods if we impute the missing miRNA expressions with their median across the 18 patients. For Scheme (c) alone, the total number of detections rose from fourteen to twenty-seven after imputation by sample median, as shown in Table S4.2 in the Supplement. ^2^ Among the sixteen miRNAs listed in Table 1, the nine miRNAs miR-1, miR-486-3p, miR-21-5p, miR-147b, miR-99a-3p, miR-455-3p, miR-1247-5p, miR-455-5p, and miR-486-5p were undetected by De Sarkar et al. (2014). For ease of reference, we will refer to these nine as “novel” miRNAs. Their relevance will be discussed in detail in Section 3.3. As mehtioned previously, De Sarkar et al. (2014) found that established cancer biomarkers hsa-miR-1 and miR-21-5p were downregulated and upregulated, respectively, in many samples, although neither was detected as significantly deregulated in their analysis (Heatmap 3). In particular, miR-1’s *p*-value was just above the BH-adjusted significance threshold in their study. Every group-adaptive BH method detected miR-1 to be significantly downregulated, regardless of the grouping scheme. TST-GBH detected miR-21-5p as significantly deregulated under all grouping schemes except the coarsest, Scheme (a). In a cluster analysis, De Sarkar et al. (2014) identified a subgroup of 13 patients with 30 deregulated miRNAs in cancer tissues (see Table 6 in De Sarkar et al., 2014). Except miR-486-3p/5p and miR-455-3p/5p, all novel miRNAs were members of this set of 30 miRNAs.

#### Overlap with prior OSCC studies

Most of the significantly deregulated miRNAs reported in Table 1, except miR-1293, miR-1247-5p, and miR-147b, have been mentioned in prior OSCC studies. Notably, hsa-miR-21, miR-31, and miR-204 are among the most frequently reported diagnostic and prognostic markers across OSCC studies (Troiano et al., 2018; Rajan et al., 2021; Ghosh et al., 2020; Min et al., 2015; Wong et al., 2016). In addition, miR-1 and miR-455 have been highlighted in surveys and meta-analyses for their prognostic relevance (Troiano et al., 2018; Ghosh et al., 2020). Importantly, miR-21, miR-204, miR-1, and miR-455 were identified only by the group-adaptive BH methods in our analysis (Table 1).

At the same time, our methods did not detect several miRNAs frequently cited for their potential prognostic value in OSCC, including miR-125b, miR-155, miR-16, miR-196a, miR-20a, and miR-32 (Min et al., 2015; Ghosh et al., 2020; Troiano et al., 2018). There are several possible reasons for this lack of overlap. First, most studies included in the cited reviews focused on tongue and floor-of-mouth cancers, which differ in clinical behavior and molecular features from gingivo-buccal squamous cell carcinoma (GBSCC), the subtype predominant in the South Asian tobacco-chewing population (Pansare et al., 2019). In fact, many of these miRNAs, e.g., miR-196a, miR-20a, and miR-32, were also absent in the dataset of De Sarkar et al. (2014). Second, the lack of overlap may also be due to the diminished statistical power, owing to the limited sample size of our data. A deeper investigation into the sources of this discrepancy is beyond the scope of this study. What is relevant here is that the group-adaptive BH methods succeeded in enhancing detections relative to BH. Nonetheless, statistical methods can not recover signals that are not present in the data.

#### Effect of data missingness

Although we restricted our analysis to complete cases, missingness may still have influenced the performance of the GBH methods. Among the miRNAs in Table 1, miR-1293 showed the highest level of missingness (50%), and evidence for its cancer-promoting role in the literature is limited. Table 1 includes five other miRNAS, miR-31-3p, miR-204-5p, miR-7-5p, miR-147b, and miR-1247-5p, with lower level of missingness (10%). Among these, the miRNAs with lower p-values, e.g., miR-31-3p, miR-204-5p, and miR-7-5p, were previously implicated in OSCC (see De Sarkar et al., 2014, for details). By contrast, evidence supporting the observed deregulation of miR-147b and miR-1247-5p, which had relatively higher p-values, is relatively limited in OSCC; we return to these miRNAs in Section 3.3. Notably, miR-1293, miR-147b, and miR-1247-5p were mostly detected by TST-GBH (Table 1, Supplementary Table S4.1). By contrast, SABHA generally did not detect any miRNAs with missing data to be significantly deregulated under the considered grouping schemes. The only exception was miR-31-3p, which was also identified by BH and is a well-established oncomiR in OSCC (Lin et al., 2022). Of the six detected miRNAs with missing values, miR-31-3p, miR-204-5p, miR-7-5p, and miR-1293 were also reported to be significantly deregulated in De Sarkar et al. (2014)’s analysis, which, we remind readers, used imputed data. We suspect that the small sample size amplified the adverse effects of missingness.

### 3.2 Control study: random group assignment

To see if our position-based groups indeed helped with detection, as a small control study, we compared our results to the case where the miRNAs were assigned randomly to the groups. To this end, we restricted our attention to the groups in Scheme (c), where the groups are based on shared chromosome number, strands, and arms. Under random assignment, the groups remained identical to those of Scheme (c) in number and size, but the miRNAs were assigned to them randomly, not based on their position. Therefore, the groups were no longer biologically informative. The random assignment of the miRNAs to groups did not break the diseased-normal association, but broke the intra-group associations. The random assignments were carried out 1000 times, resulting in 1000 replications. The group-adaptive BH methods were performed in each replication. The BH was not performed because it has no dependence on group assignments.

### 3.2.1 Effect of random grouping on detections

With the exception of miR-135b-3p, the miRNAs frequently detected under random assignments (in more than 50% of replications) were the same ones identified under Scheme (c) as significantly deregulated by at least one method. Table 2 provides the detection frequencies of these miRNAs. Table 3 provides summary statistics on the detections of each method across the 1000 replications, and report their overlap with the three BH-detected miRNAs: miR-133a-3p, miR-31-3p, and miR-206. On average, the number of detections under random group assignments dropped to three for both SABHA and LSL-GBH. Moreover, most detections by LSL-GBH (about two out of three) and all detections by SABHA overlapped with the three BH miRNAs (Table 3). Table 1 shows that SABHA and LSL-GBH collectively detected cancer biomarkers such as miR-31-5p, miR-1, and the oncomiR miR-7-5p to be significantly deregulated under the original, non-randomized, grouping Scheme (c) (Lu et al., 2019; Safa et al., 2020; Jung et al., 2012). However, LSL-GBH and SABHA did not detect these miRNAs in more than two-third of the random assignments, respectively (Table 2). Therefore, LSL-GBH and SABHA’s detection of these miRNAs under Scheme (c) may be attributable to the additional power gained from position-based grouping.

**Table 2.** Detection of significantly deregulated miRNAs under 1000 random reassignments of Scheme (c). Each replication randomly reassigns miRNAs to groups in Scheme (c), preserving group sizes and diseased–normal pairing but breaking intra-group associations. This table lists the miRNAs that were either (1) detected to be significantly deregulated under original (non-randomized) Scheme (c) by at least one of the four FDR-control methods or (2) identified as significantly deregulated in more than 50% of the random group assignments by at least one group-adaptive BH method. The percentages under TST (stands for TST-GBH), LSL (stands for LSL-GBH), and SABHA represent the proportion of 1,000 random assignments in which each miRNA was identified as significantly deregulated by the corresponding method. A cross x in these columns indicates that the miRNA was detected under the original (non-randomized) Scheme (c). Only hsa-miR-135b-3p (highlighted) was not detected under the original Scheme (c). The “Chr.” column gives the chromosome, the “Str.” column indicates the chromosome strand, and the “arm” column specifies the arm. The “ΔΔCt” column provides the average ΔΔCt value for each miRNA across 18 patients. The “p-value” column shows raw p-values from paired t-tests. Rows are sorted by p-values, and the arrows following them denote whether the expression was upregulated (↑) or downregulated (↓) according to the sign of ΔΔCt. The “Missing pairs” column provides the number of missing pairs of observations for each miRNA. *These miRNAs were reported to be significantly deregulated by De Sarkar et al. (2014). † These miRNAs were detected by BH to be significantly deregulated.

| miRNA | Chr. | Str. | Arm | $\Delta\Delta Ct$ | p-value | Miss<br>-ing<br>pairs | TST | LSL | SABHA |
| --- | --- | --- | --- | --- | --- | --- | --- | --- | --- |
| hsa-miR-133a-3p<sup>*†</sup> | 18 | - | q | 6.7 | $5.32E^{-05}$ ↓ | 0 | x (100%) | x (56%) | x (100%) |
| hsa-miR-31-3p<sup>*†</sup> | 9 | - | p | -3.8 | $1.31E^{-04}$ ↑ | 2 | x (100%) | x (51%) | x (100%) |
| hsa-miR-206<sup>*†</sup> | 6 | + | p | 6.0 | $1.97E^{-04}$ ↓ | 0 | x (100%) | 48% | x (100%) |
| hsa-miR-31-5p<sup>*</sup> | 9 | - | q | -3.4 | $6.18E^{-04}$ ↑ | 0 | x (100%) | x (33%) | x (3%) |
| hsa-miR-204-5p<sup>*</sup> | 9 | - | q | 4.6 | $8.81E^{-04}$ ↓ | 2 | x (94%) | 27% | 3% |
| hsa-miR-1 | 18 | - | q | 5.2 | $9.38E^{-04}$ ↓ | 0 | x (95%) | x (27%) | x (3%) |
| hsa-miR-7-5p<sup>*</sup> | 15 | + | q | -3.1 | $9.50E^{-04}$ ↑ | 2 | x (94%) | x (27%) | 3% |
| hsa-miR-1293<sup>*</sup> | 12 | - | q | -4.8 | $2.77E^{-03}$ ↑ | 9 | x (76%) | 9% | 0% |
| hsa-miR-486-3p | 8 | - | p | 2.4 | $2.81E^{-03}$ ↓ | 0 | x (73%) | 8% | 0% |
| hsa-miR-21-5p | 17 | + | q | -2.2 | $3.67E^{-03}$ ↑ | 0 | x (71%) | 6% | 0% |
| hsa-miR-147b | 15 | + | q | -2.1 | $5.52E^{-03}$ ↑ | 2 | x (51%) | 3% | 0% |
| hsa-miR-135b-3p | 1 | - | q | -2.0 | $5.73E^{-03}$ ↑ | 2 | 50% | 3% | 0% |
| hsa-miR-99a-3p | 21 | + | q | 2.8 | $5.83E^{-03}$ ↓ | 0 | x (49%) | 3% | 0% |
| hsa-miR-1247-5p | 14 | - | q | 2.4 | $8.93E^{-03}$ ↓ | 2 | x (31%) | 1% | 0% |
| hsa-miR-486-5p | 8 | - | p | 2.0 | $1.78E^{-02}$ ↓ | 0 | x (8%) | 0% | 0% |

**Table 3.**
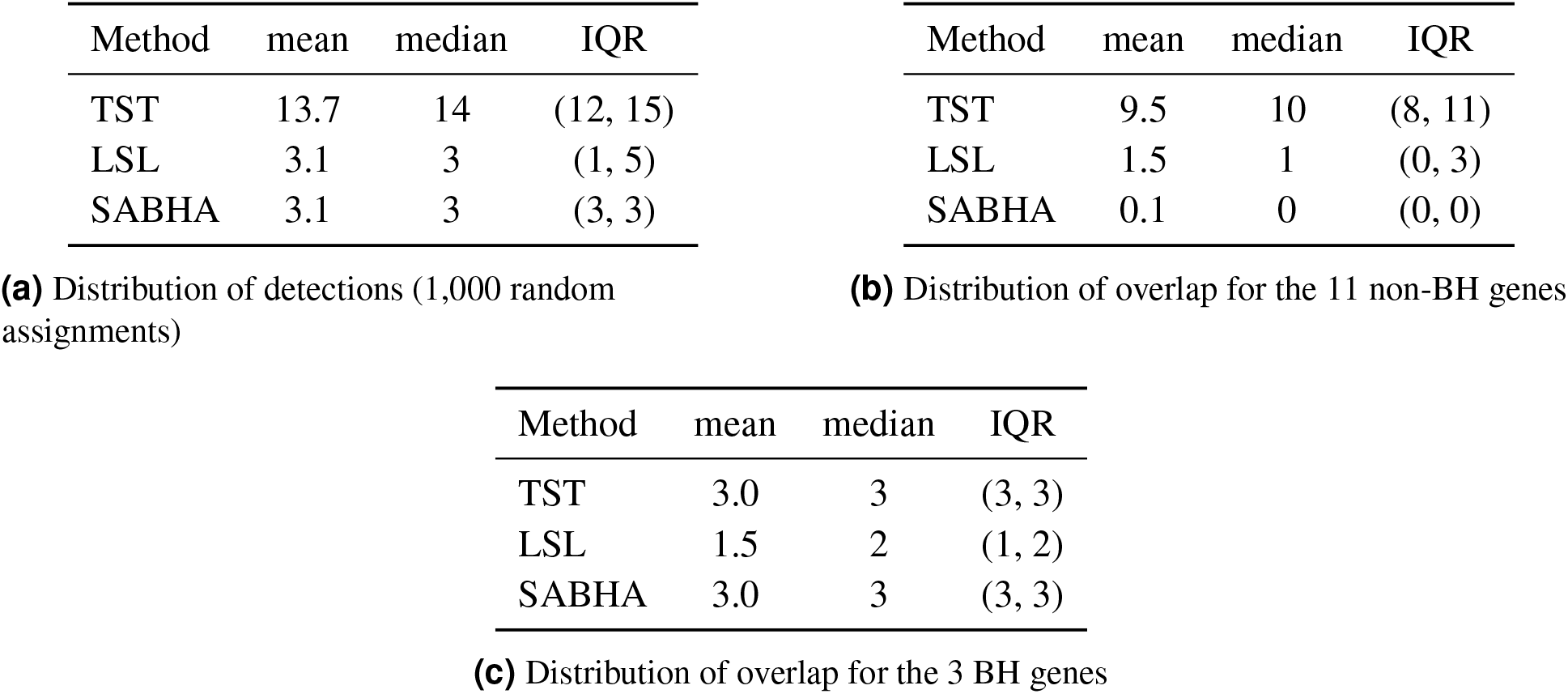
Summary statistics of significantly deregulated miRNAs from 1,000 random assignments. Each replication randomly reassigns miRNAs to groups in Scheme (c), preserving group sizes and diseased–normal pairing but breaking intra-group associations. The summary statistics are reported for group-adaptive methods TST-GBH (TST), LSL-GBH (LSL), and SABHA, listed under the column “Method.” (i) This table shows mean, median, and interquartile range (IQR) of the number of significantly deregulated miRNAs across 1,000 random assignments; (ii) This table shows mean, median, and IQR of the number of detected miRNAs that overlaps with the three BH-detected miRNAs (hsa-miR-133a-3p, miR-31-3p, and miR-206); and (iii) This table shows mean, median, and IQR of the number of detected miRNAs that overlaps with the non-BH miRNAs. The non-BH miRNAs are the 11 miRNAs detected under Scheme (c) by group-adaptive BH methods but not by BH (see Table 1). In all tables, the summary statistics are calculated across the 1,000 replications.

In contrast to LSL-GBH and SABHA, the average number of detections by TST-GBH did not decline substantially under random group assignment. On average, TST-GBH still identified 14 miRNAs as significantly deregulated (Table 3(i)). These detections always included the three miRNAs found by BH, along with miR-31-5p (Table 2). Among the remaining detections, about six miRNAs overlapped with those identified by this method under Scheme (c), while the rest varied randomly across samples. This pattern suggested that position-based grouping may have helped TST-GBH detect certain miRNAs, but the effect is less prominent than for SABHA and LSL-GBH. Overall, the control study provided stronger supporting evidence for SABHA and LSL-GBH in this dataset.

As mentioned earlier, miR-135b-3p (upregulated with ΔΔ*Ct* = ™2.04) is the only new miRNA frequently detected across the replications. It is identified as significantly deregulated by TST-GBH in 50% of the replications (Table 2). It was also detected by TST-GBH under the finer grouping schemes derived from Scheme (c) when the maximum group size dropped below 6. miR-135b is frequently reported as upregulated in several cancers, including OSCC, and has been speculated to act as a promoter of OSCC (Olasz et al., 2015; Shao et al., 2025). Apart from miR-135b-3p, 43 additional miRNAs were detected in at least one of the 1,000 replications. However, no method detected them to be significantly deregulated in more than 40% of the replications.

### 3.3 Relevance of the novel miRNAs

Although it would have been desirable to experimentally validate the pathways downstream of the novel miRNAs, such analyses were beyond the scope of the current study because of the exhaustion of biospecimens used by De Sarkar et al. (2014). However, several other functional works provided supporting evidence for the role of the “novel” miRNAs in carcinogenesis. Since 10 of the 12 cancer tissues assessed by Singh et al. (2017) overlap with the cohort studied in De Sarkar et al. (2014), their whole transcriptome analysis allowed for correlative analysis. We further compared our results with previously published onco-miRNAome studies.

Singh et al. (2017) assessed the differential expression of the genes (measured as mRNA transcripts) using EdgeR. EdgeR uses the BH method as a default choice for multiple-testing correction. A gene was considered to be significantly deregulated if it showed at least 1.5-fold change and an FDR-adjusted p-value *<* 0.05. Similar to us, they avoided imputations. The paper reports that genes with low or inconsistent expression values were removed before analysis. Eight of the nine novel miRNAs had some of their known target genes significantly deregulated in the opposite direction of the miRNA’s deregulation pattern in Singh et al. (2017)’s analysis. Table 4 lists these genes with their average fold changes, along with source articles reporting these genes as targets of the corresponding miRNAs.

**Table 4.**
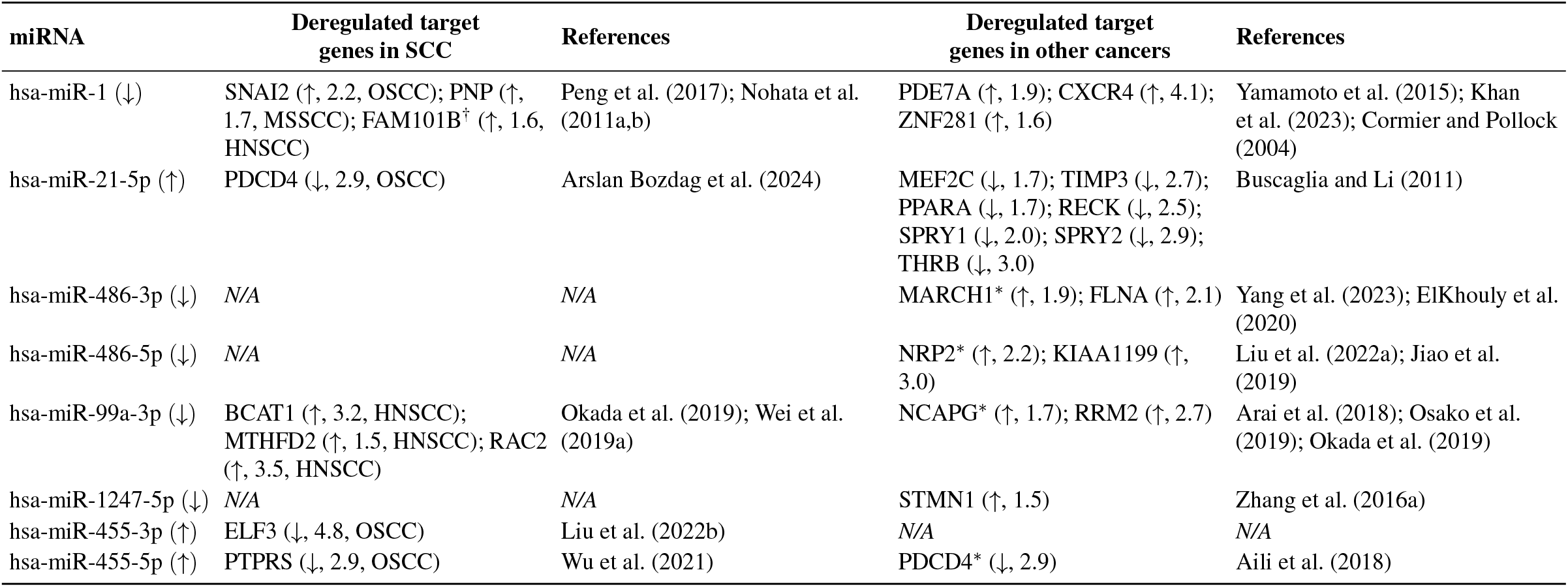
Target genes of the newly discovered miRNAs with significantly deregulated expressions, as identified by Singh et al. (2017). All genes in the “Deregulated target genes in SCC” column, except FAM101B, are validated targets of the corresponding miRNA in the squamous cell carcinoma (SCC) subtype indicated next to each gene. Source references are provided in the adjacent “ References ” column. In addition, Singh et al. also reported genes that have been described as miRNA targets in other cancers but not specifically in SCC. These are listed in a separate column with their corresponding references in the adjacent “References” column. All genes reported in this table were significantly deregulated in Singh et al. (2017)’s whole transcriptome analysis in the opposite direction of the miRNA deregulation pattern. For each gene, the average fold change and direction of deregulation are shown in parentheses next to the gene name. The deregulation patterns of the corresponding miRNAs are also provided. **Abbreviations:** SCC: squamous cell carcinoma; MSSCC: maxillary sinus SCC; HNSCC: head and neck SCC. ^*^ Associated with OSCC but, to our knowledge, not validated as direct targets in OSCC (see Section 3.3). ^†^ Nohata et al. (2011b) found that FAM101B was significantly downregulated in HNSCC cells following miR-1 transfection and contained a predicted miR-1 target site. However, to our knowledge, FAM101B has not been validated as a direct target of miR-1.

miR-147b was the only miRNA without any significantly deregulated target gene in Singh et al. (2017)’s analysis. To our knowledge, miR-147b has not been previously characterized in OSCC. However, it has been implicated as an oncogenic miRNA in other cancers, such as lung cancer, which is consistent with its observed upregulation in our study (Zhang et al., 2019; Wang et al., 2020; Meng et al., 2013; Cui et al., 2015). Among the remaining novel miRNAs, miR-1, miR-486-3p/5p, miR-1247-5p, and miR-99a-3p were significantly downregulated in our analysis, where miR-21-5p and miR-455-3p/5p were significantly upregulated. These patterns of deregulation are not only consistent with Singh et al. (2017)’s analysis, but also with prior knowledge, which we detail in the following discussion.

#### miR-1 (downregulated) miR-1

was downregulated in De Sarkar et al. (2014)’s data, which is consistent with its established roles as a tumor suppressor (Safa et al., 2020). The reduced expression of miR-1 in OSCC has been associated with advanced disease and poorer prognosis (Peng et al., 2017). In OSCC, miR-1 suppresses epithelial–mesenchymal transition (EMT) by directly targeting the transcription factor SNAI2 or Slug (Peng et al., 2017). SNAI2’s expression was significantly upregulated with an average fold change of 2.2 in Singh et al. (2017)’s analysis. Nohata et al. (2011a) reported that miR-1 targets the oncogene PNP in maxillary sinus squamous cell carcinoma (MSSCC), a subtype of SCC. Its upregulation with an average fold change of 1.7 in Singh et al. (2017) suggests the possibility that the PNP–miR-1 pathway may play a similar role in OSCC. Additional target genes of miR-1 were found with significantly upregulated expression in Singh et al. (2017)’s analysis, including well-established oncogenes such as PDE7A in endometrial cancer (Yamamoto et al., 2015), CXCR4 in small cell lung cancer (Khan et al., 2023) and thyroid cancer (Leone et al., 2011), and ZNF281 in soft tissue sarcomas (Cormier and Pollock, 2004) See Table 4 for more details.

#### miR-21-5p (upregulated)

miR-21-5p is a ubiquitous oncogene, widely proposed as a prognostic biomarker for OSCC, and the most frequently overexpressed miRNA in many cancers (Schneider et al., 2018; Dioguardi et al., 2022; Troiano et al., 2018). Its overexpression in our analysis is therefore consistent with its well-documented oncogenic role in OSCC (Feng and Tsao, 2016; Reis et al., 2010; Reddy et al., 2024; Schneider et al., 2018).

PDCD4, a target gene of miR-21-5p, associated with progression and metastasis in OSCC (cf. Reis et al., 2010; Arslan Bozdag et al., 2024), was significantly downregulated with an average fold change of 2.9 in Singh et al. (2017). PDCD4 is regarded as one of the principal targets of miR-21 (Jenike and Halushka, 2021). Table 4 shows that seven additional target genes of miR-21, namely MEF2C, TIMP3, PPARA, RECK, SPRY1, SPRY2, and THRB, were also significantly downregulated in Singh et al. (2017). According to Buscaglia and Li (2011), miR-21 suppresses the activities of these genes, whose downregulation is linked to multiple hallmarks of cancer (Hanahan and Weinberg, 2011).

Notably, using OSCC samples from Indian patients, Manikandan et al. (2016) compared miRNA expression profiles between moderately and well-differentiated tumor cells relative to normal squamous epithelium. Well-differentiated tumors are generally more aggressive than moderately differentiated tumors. As shown in the heatmap in Figure S4.2 of the Supplement, hsa-miR-21-5p expression was generally higher in the patients with more aggressive tumors (average fold change 1.93), indicating that miR-21 may play a role in early-stage tumorigenesis in OSCC.

#### miR-486-3p/5p (downregulated)

miR-486-3p exhibits tumor suppressing activities in many cancers including OSCC, where its downregulation is associated with migration and invasion (ElKhouly et al., 2020; Yang et al., 2020). miR-486-5p’s role in OSCC is more controversial but recent studies support its tumor suppressing activity in OSCC, which aligns with the observed downregulation (Yan et al., 2016; Soga et al., 2013). Among the known target genes of miR-486, FLNA, MARCH1, KIAA1199, and NRP2 were significantly upregulated in Singh et al. (2017). FLNA and MARCH1 have been validated as miR-486-3p targets in laryngeal squamous cell carcinoma (ElKhouly et al., 2020) and osteosarcoma (Yang et al., 2023), respectively, whereas KIAA1199 and NRP2 were confirmed as direct miR-486-5p targets in papillary thyroid cancer (Jiao et al., 2019) and colorectal cancer (Liu et al., 2019), respectively. Among these, MARCH1 and NRP2 have been linked to prognosis in OSCC. MARCH1 modulates cell proliferation and apoptosis (Liu et al., 2022a), while NRP2 influences proliferation, migration, and invasion in OSCC (Kang et al., 2021). However, known target genes of miR-486-3p in OSCC, e.g., DDR1, ANK1, FGF44, etc., (cf. ElKhouly et al., 2020, for details) were absent from Singh et al. (2017)’s list of significantly deregulated genes.

#### miR-99a-3p (downregulated)

miR-99a-3p has been reported to function as a tumor suppressor in several cancers, including OSCC (Chen et al., 2018; Wang et al., 2025). To our knowledge, literature on validated targets of this miRNA in OSCC is limited. However, CDK6, a validated target of miR-99b-3p in OSCC (cf. Wang et al., 2025), was found to be upregulated in Singh et al. (2017) with an average fold change of 1.7. The cancer biomarker NCAPG, identified as a direct target of miR-99a-3p in prostate cancer (Arai et al., 2018), has been shown to regulate cell proliferation, cell cycle progression, and apoptosis in OSCC (Jianing et al., 2021). In Singh et al. (2017), NCAPG was also upregulated with an average fold change of 1.7, raising the possibility that it may also be a target of miR-99a-3p in OSCC. More broadly, Okada et al. (2019); Wei et al. (2019a) examined the tumor suppressor roles of miR-99a-3p in head and neck squamous cell carcinoma and reported several direct targets. Among these, BCAT1, MTHFD2, and RAC2 were significantly upregulated in Singh et al. (2017)’s analysis. In addition, RRM2, a direct target of miR-99a-3p in renal carcinoma (cf. Osako et al., 2019), was also found to be upregulated with average fold change of 2.7 in that study.

#### miR-1247-5p (downregulated)

miR-1247-5p typically acts as a tumor suppressor in several cancers, which is consistent with our observations (Chu et al., 2017; Yi et al., 2017; Zhang et al., 2016a; Wei et al., 2019b). Although research on the role of miR-1247 in OSCC is limited (Liu et al., 2020), it has been found to be downregulated in cutaneous squamous cell carcinoma, a subtype of SCC (An et al., 2019). Moreover, the oncogene STMN1, reported as a functional target of miR-1247 in non-small-cell lung cancer (Zhang et al., 2016b), was significantly upregulated (average fold change 1.5) in the analysis of Singh et al. (2017).

#### miR-455-3p (upregulated)

MiR-455-3p has been studied in several cancers, and its role is context-dependent. Broadly, it has been reported to act both as a tumor suppressor and an oncogene, depending on the tissue context (Ye et al., 2023). Some recent studies have highlighted its oncogenic roles in OSCC, in line with our observations (Li et al., 2020; Liu et al., 2022b). Liu et al. (2022b) showed that miR-455-3p targets the tumor suppressor transcription factor ELF3 in OSCC, which was significantly downregulated (average fold change 4.8) in Singh et al. (2017)’s analysis. miR-455-3p’s oncogenic activity and elevated expression have also been reported in other carcinomas, including esophageal squamous cell carcinoma (Liu et al., 2017), colorectal cancer (Ye et al., 2023), and triple-negative breast cancer (Li et al., 2016).

#### vmiR-455-5p (upregulated)

miR-455-5p is suspected to promote cancer cell migration and invasion in OSCC, which aligns with the observed upregulation (Hsiao et al., 2023; Cheng et al., 2016). Wu et al. (2021) suggested that miR-455-5p modulates cell proliferation and growth in OSCC by targeting PTPRS. PTPRS was significantly downregulated with average fold change 2.9 in Singh et al. (2017). This miRNA also exhibits oncogenic activities in breast cancer (Aili et al., 2018), lung cancer (Wang et al., 2017), as well as bladder urothelial carcinoma (Hamilton et al., 2013). Aili et al. (2018) showed that miR-455-5p targets the tumor suppressor gene PDCD4 in breast cancer, which was significantly downregulated with an average fold change of 2.9 in Singh et al. (2017). PDCD4, also a primary target of miR-21-5p, is associated with nodal metastasis and invasion in OSCC (Reis et al., 2010).

## DISCUSSION

Biological studies often involve testing many hypotheses simultaneously, increasing the risk of false discoveries. Controlling the FDR allows investigators to strike a balance between the detection of true positive findings and minimizing the likelihood of false discoveries. The BH method is arguably the most widely used approach for FDR control in bioinformatics (Haynes, 2013). However, it compromises statistical power because it remains agnostic to additional structure among the hypotheses (Hu et al., 2010; Li and Barber, 2019). This limitation is especially problematic in small-sample studies where the number of hypotheses far exceeds the sample size, such as the oral cancer miRNA dataset reanalyzed here. In the dataset’s original report, De Sarkar et al. (2014) observed that OSCC-related miRNAs such as miR-1 and miR-21 failed to meet the FDR cutoff with the stringent BH method, although exploratory analyses suggested their deregulation. Moreover, a follow-up study by Singh et al. (2017) found that several target genes of these miRNAs were significantly deregulated in a cohort largely overlapping with De Sarkar et al. (2014)’s.

Although various types of structural information can be leveraged in the FDR control step, here we focused specifically on group structure. Group-adaptive BH methods are modifications of the standard BH method that use information on group structure to improve statistical power. Because De Sarkar et al. (2014)’s data were generated under rigorous analysis techniques, we used it as a test case to examine the potential advantages of these methods over BH. In this proof-of-principle study, our aim was not to identify new biomarkers, but to illustrate how incorporating simple biological structure, here, positional groupings of miRNAs, can influence the outcome of FDR control.

This study considered three group-adaptive BH methods, LSL-GBH, TST-GBH, and SABHA, chosen for their simplicity, feasibility in small-sample settings, and suitability for demonstrating how group information can be incorporated into the FDR control step. We compared several positional grouping schemes, in which miRNAs were clustered by chromosomal location with increasing levels of refinement. Across methods, the number of detections generally tended to increase as grouping schemes became finer. SABHA was the most conservative and showed the least sensitivity to changes in grouping schemes. In contrast, TST-GBH was the most liberal and most sensitive to changes in group sizes, consistent with earlier reports of its behavior (Sarkar and Zhao, 2022). In the final results, we considered grouping schemes with moderately sized groups, where all methods were relatively stable.

The group-adaptive BH methods collectively detected several miRNAs that were not detected as significantly deregulated by BH, including miR-1 and miR-21. Most of these miRNAs have been previously associated with OSCC with deregulation patterns consistent with our study (Section 3.3). These miRNAs had several targets deregulated in the opposite direction (of the miRNA expression) in the follow-up transcriptome analysis of Singh et al. (2017), providing further support for their biological relevance. Despite TST-GBH being more liberal, its list of detections contained only two miRNAs without prior implications in OSCC, namely, miR-147b and miR-1293. Both miRNAs, especially miR-1293, had relatively high level of missingness. By contrast, all detections of LSL-GBH and SABHA, including those with missing values, had previous implications in OSCC. Taken together, in this dataset, group-adaptive BH methods showed potential for detecting signals missed by BH, although sensitivity to grouping and missingness varied across methods.

We recognize that grouping of miRNAs based on broad spatial information is overly simplistic. Groupings could also be defined by other forms of functional or genetic relatedness, such as ancestral similarity. However, exploring these alternatives was beyond the scope of this proof-of-principle analysis. While positional information does not capture all dimensions of functional relatedness, it provides an interpretable grouping strategy appropriate for the limited sample size of our dataset. Our control study with random group assignments suggested that the additional detections by LSL-GBH and SABHA may be attributable to the incorporation of positional grouping, indicating that even basic structural information has the potential to improve detections.

Alternative grouping strategies, such as ontology-based assignments (Liu et al., 2015) or data-driven clustering methods, e.g., k-means, tight clustering (Karmakar et al., 2019), WGCNA (Horvath, 2011), may also be useful and could provide further gains in statistical power. The data-dependent clustering approaches, however, must be applied with caution, as recent work has shown that reusing the same data for both clustering and hypothesis testing can inflate false discoveries (Lähnemann et al., 2020; Gao et al., 2020b). We remind the readers that independence between groups is not required. Group-adaptive BH methods are generally most advantageous when most hypotheses are null and the intra-group association is higher than inter-group association (Hu et al., 2010; Chu et al., 2017). Existing literature suggests the first condition is likely in miRNAome data (Osan et al., 2021), and our analyses in Section 2.4.1 indicated that the second may also plausibly hold for our location-based grouping schemes.

In addition to the group-adaptive BH methods studied here, other approaches exist for incorporating group structure into FDR control, such as independent hypothesis weighting (Ignatiadis et al., 2016), q-values (Storey, 2002), and knock-off (Sesia et al., 2021b). Our intent was not to benchmark all available approaches or to claim superiority of the group-adaptive BH methods. Rather, in this proof-of-concept analysis, we aimed to demonstrate that even simple methods that exploit external grouping information can make discoveries that are missed by the standard BH procedure. The promising performance observed here suggests that more sophisticated approaches may also be effective, but this remains to be evaluated systematically in future work. The choice of method should ultimately depend on the scale and characteristics of the dataset.

Although our analysis was based on a miRNAome dataset, the strategy of incorporating group information into the FDR control step can be applied to any multiple testing scenario. Of course, controlled future analyses are needed to test such generalizability. In particular, extrapolation to larger datasets generated by high-throughput sequencing technologies remains to be tested. Of particular interest are the datasets pertaining to treatment-emerging cancer subtypes, which often have small sample sizes similar to our study (Aggarwal et al., 2019; Clermont et al., 2019). These types of cancers arise as a result of treatment-related factors, potentially as a consequence of the treatment itself or because of the underlying conditions being treated. The genes responsible for such cancer subtypes are still not fully elucidated. Future studies aiming to unveil these genetic factors will necessitate FDR control because of multiple hypothesis testing.

## CONCLUSION

The conservativeness of the BH method can limit scientific discoveries, especially in studies where the sample size is small but hunderds of hypotheses need to be tested simultaneously. Using the miRNAome dataset of De Sarkar et al. (2014), we demonstrated that recent modifications of the BH method that incorporate group structure can offer advantages in such settings. Incorporating even simple group information, such as positional grouping among miRNAs, appeared to contribute to improved detections, although sensitivity to grouping varied across methods. While our study was limited to a single moderately scaled dataset, the results highlight the potential of group-adaptive BH methods for small-sample studies where statistical power is a concern. Future work should assess these methods across datasets of varying scales and designs, and explore alternative grouping strategies and more advanced FDR control procedures to establish their broader applicability.

## ACKNOWLEDGMENTS

The authors are thankful to Editage for providing editing support to an earlier version of the manuscript.

## Supplement

Attached, contains more details on miRNAome data generation, missingness, and additonal tables and plots.

## Code

Code is made available on Github (Koner et al., 2023).

## 5. APPENDIX

### 5.1 Simple random effect model to quantify the inter-group and intra-group correlation

To formalize the setup, we will introduce some new notation. Denote by {*Y*_*i j*_: *j* = 1, …, 522} the ΔΔ*Ct* values of the *j*th miRNA of the *i*th subject/ patient. Let *g*(*j*) be the group indicator of the *j*th miRNA, e.g., if the *j*th miRNA belongs to the first group, then *g*(*j*) = 1. The group index *g*(*j*) takes value between 1 and 67.

We will posit a random effect model Searle et al. (2009) on the ΔΔ*Ct* values as

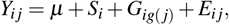

where *µ* is the overall mean, *S*_*i*_ is the random effect of the ith patient, *G*_*ig*(*j*)_ is the random effect of the *g*(*j*)th group for the *i*th patient, and *E*_*i j*_ is the random error for the *i*th subject and the *j*th miRNA. The random error can represent independent measurement errors or other underlying biological factors. We let the collections of random variables {*S*_*i*_: *i* = 1, …, 18}, {*G*_*ig*(*j*)_: 1 ≤ *i* ≤ 18, 1 ≤ *j* ≤ 522}, and {*E*_*i j*_: 1 ≤ *i, j* ≤ 522} to be independent of each other. We will also assume that the *S*_*i*_’s are identically distributed as 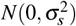, the *G*_*ig*(*j*)_’s are identically distributed as 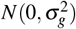, and the *E*_*i j*_’s are identically distributed as 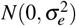 where 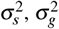, and 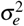 are unknown positive numbers.

Under the above model, the correlation between the ΔΔ*Ct* values of two miRNAs belonging to two different groups within the same subject is constant. The above correlation will thus be a natural candidatefor the inter-group correlation. Straight-forward algebra shows that the inter-group correlation equals 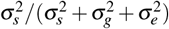 Likewise, under our model, the ΔΔ*Ct* values of two miRNAs belonging to the same group within the same subject have identical correlations across the subjects and the miRNAs – this correlation will be a natural candidate for the intra-group correlation. The intra-group correlation is given by 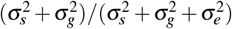. We used the R package `lme4` to fit the model and obtain estimates of the intra- and inter-group correlations.

## Footnotes

1 SABHA can adapt to a wide range of structures, including, but not limited to, grouped hypotheses (Li and Barber, 2019).

2 Table S4.2 shows that some of the additional detections in the median-imputed data had no missing data. This is unsurprising because all four FDR-control methods rely on BH-type ranking of p-values to determine the detection threshold. Imputation-led p-value deflation can make the overall p-value threshold less stringent in such cases, as discussed in Section 2.1.1.

## REFERENCES

Aggarwal, R. R., Quigley, D. A., Huang, J., Zhang, L., Beer, T. M., Rettig, M. B., Reiter, R. E., Gleave, M. E., Thomas, G. V., Foye, A., et al. (2019). Whole-genome and transcriptional analysis of treatment-emergent small-cell neuroendocrine prostate cancer demonstrates intraclass heterogeneity. Molecular Cancer Research, 17(6):1235–1240.

Aili, T., Paizula, X., and Ayoufu, A. (2018). miR-455-5p promotes cell invasion and migration in breast cancer. Molecular Medicine Reports, 17(1):1825–1832.

An, X., Liu, X., Ma, G., and Li, C. (2019). Upregulated circular rna circ 0070934 facilitates cutaneous squamous cell carcinoma cell growth and invasion by sponging mir-1238 and mir-1247–5p. Biochemical and Biophysical Research Communications, 513(2):380–385.

Arai, T., Okato, A., Yamada, Y., Sugawara, S., Kurozumi, A., Kojima, S., Yamazaki, K., Naya, Y., Ichikawa, T., and Seki, N. (2018). Regulation of ncapg by mir-99a-3p (passenger strand) inhibits cancer cell aggressiveness and is involved in crpc. Cancer medicine, 7(5):1988–2002.

Arslan Bozdag, L., Açik, L., Ersoy, H. E., Bayir, Ö., Korkmaz, M. H., Mollaoglu, N., and Gultekin, S. E. (2024). Pdcd 4 and mir-21 are promising biomarkers in the follow-up of oed in liquid biopsies. Oral Diseases, 30(6):3873–3883.

Benjamini, Y. and Heller, R. (2007). False discovery rates for spatial signals. Journal of the American Statistical Association, 102(480):1272–1281.

Benjamini, Y. and Hochberg, Y. (1995). Controlling the false discovery rate: a practical and powerful approach to multiple testing. Journal of the Royal statistical society: series B (Methodological), 57(1):289–300.

Benjamini, Y. and Hochberg, Y. (2000). On the adaptive control of the false discovery rate in multiple testing with independent statistics. Journal of educational and Behavioral Statistics, 25(1):60–83.

Benjamini, Y., Krieger, A. M., and Yekutieli, D. (2006). Adaptive linear step-up procedures that control the false discovery rate. Biometrika, 93(3):491–507.

Benjamini, Y. and Yekutieli, D. (2001). The control of the false discovery rate in multiple testing under dependency. Annals of statistics, pages 1165–1188.

Buscaglia, L. E. B. and Li, Y. (2011). Apoptosis and the target genes of microrna-21. Chinese journal of cancer, 30(6):371.

Bustin, S. A., Benes, V., Garson, J. A., Hellemans, J., Huggett, J., Kubista, M., Mueller, R., Nolan, T., Pfaffl, M. W., Shipley, G. L., et al. (2009). The miqe guidelines: Minimum i nformation for publication of q uantitative real-time pcr e xperiments.

Chen, L., Hu, J., Pan, L., Yin, X., Wang, Q., and Chen, H. (2018). Diagnostic and prognostic value of serum miR-99a expression in oral squamous cell carcinoma. Cancer Biomarkers, 23(3):333–339.

Cheng, C.-M., Shiah, S.-G., Huang, C.-C., Hsiao, J.-R., and Chang, J.-Y. (2016). Up-regulation of miR-455-5p by the TGF-β –SMAD signalling axis promotes the proliferation of oral squamous cancer cells by targeting ube2b. The Journal of pathology, 240(1):38–49.

Chu, Y., Fan, W., Guo, W., Zhang, Y., Wang, L., Guo, L., Duan, X., Wei, J., and Xu, G. (2017). mir-1247-5p functions as a tumor suppressor in human hepatocellular carcinoma by targeting wnt3. Oncology reports, 38(1):343–351.

Clermont, P.-L., Ci, X., Pandha, H., Wang, Y., and Crea, F. (2019). Treatment-emergent neuroendocrine prostate cancer: molecularly driven clinical guidelines. International Journal of Endocrine Oncology, 6(2):IJE20.

Cormier, J. N. and Pollock, R. E. (2004). Soft tissue sarcomas. CA: a cancer journal for clinicians, 54(2):94–109.

Cui, R., Meng, W., Sun, H.-L., Kim, T., Ye, Z., Fassan, M., Jeon, Y.-J., Li, B., Vicentini, C., Peng, Y., et al. (2015). Microrna-224 promotes tumor progression in nonsmall cell lung cancer. Proceedings of the National Academy of Sciences, 112(31):E4288–E4297.

De Sarkar, N., Roy, R., Mitra, J. K., Ghose, S., Chakraborty, A., Paul, R. R., Mukhopadhyay, I., and Roy, B. (2014). A quest for mirna bio-marker: a track back approach from gingivo buccal cancer to two different types of precancers. PloS one, 9(8):e104839.

Dioguardi, M., Spirito, F., Sovereto, D., Alovisi, M., Troiano, G., Aiuto, R., Garcovich, D., Crincoli, V., Laino, L., Cazzolla, A. P., et al. (2022). Microrna-21 expression as a prognostic biomarker in oral cancer: Systematic review and meta-analysis. International Journal of Environmental Research and Public Health, 19(6):3396.

Dudoit, S., Shaffer, J. P., and Boldrick, J. C. (2003). Multiple hypothesis testing in microarray experiments. Statistical Science, 18(1):71–103.

Efron, B. and Tibshirani, R. (2002). Empirical bayes methods and false discovery rates for microarrays. Genetic epidemiology, 23(1):70–86.

ElKhouly, A. M., Youness, R., and Gad, M. (2020). Microrna-486-5p and microrna-486-3p: Multifaceted pleiotropic mediators in oncological and non-oncological conditions. Non-coding RNA research, 5(1):11–21.

Feng, Y.-H. and Tsao, C.-J. (2016). Emerging role of microrna-21 in cancer. Biomedical reports, 5(4):395–402.

Friston, K. J. (2003). Statistical parametric mapping. Springer.

Gao, L. L., Bien, J., and Witten, D. (2020a). Selective inference for hierarchical clustering. arXiv preprint 2012.02936.

Gao, L. L., Bien, J., and Witten, D. (2020b). Selective inference for hierarchical clustering. arXiv preprint 2012.02936.

Genovese, C. R., Roeder, K., and Wasserman, L. (2006). False discovery control with p-value weighting. Biometrika, 93(3):509–524.

Ghosh, R. D., Pattatheyil, A., and Roychoudhury, S. (2020). Functional landscape of dysregulated micrornas in oral squamous cell carcinoma: clinical implications. Frontiers in oncology, 10:619.

Goeman, J. J. and Solari, A. (2014). Multiple hypothesis testing in genomics. Statistics in medicine, 33(11):1946–1978.

Hamilton, M. P., Rajapakshe, K., Hartig, S. M., Reva, B., McLellan, M. D., Kandoth, C., Ding, L., Zack, T. I., Gunaratne, P. H., Wheeler, D. A., et al. (2013). Identification of a pan-cancer oncogenic microrna superfamily anchored by a central core seed motif. Nature communications, 4(1):2730.

Hanahan, D. and Weinberg, R. A. (2011). Hallmarks of cancer: the next generation. cell, 144(5):646–674.

Haynes, W. (2013). Benjamini–hochberg method. In Encyclopedia of systems biology, pages 78–78. Springer.

Horvath, S. (2011). Weighted network analysis: applications in genomics and systems biology. Springer Science & Business Media.

Hsiao, S.-Y., Weng, S.-M., Hsiao, J.-R., Wu, Y.-Y., Wu, J.-E., Tung, C.-H., Shen, W.-L., Sun, S.-F., Huang, W.-T., Lin, C.-Y., et al. (2023). MiR-455-5p suppresses PDZK1IP1 to promote the motility of oral squamous cell carcinoma and accelerate clinical cancer invasion by regulating partial epithelial-to-mesenchymal transition. Journal of Experimental & Clinical Cancer Research, 42(1):40.

Hu, J. X., Zhao, H., and Zhou, H. H. (2010). False discovery rate control with groups. Journal of the American Statistical Association, 105(491):1215–1227.

Hughes, R. A., Heron, J., Sterne, J. A., and Tilling, K. (2019). Accounting for missing data in statistical analyses: multiple imputation is not always the answer. International journal of epidemiology, 48(4):1294–1304.

Ignatiadis, N., Klaus, B., Zaugg, J. B., and Huber, W. (2016). Data-driven hypothesis weighting increases detection power in genome-scale multiple testing. Nature methods, 13(7):577–580.

Jenike, A. E. and Halushka, M. K. (2021). mir-21: a non-specific biomarker of all maladies. Biomarker Research, 9(1):1–7.

Jianing, L., Shiqun, S., Jia, L., Xuetao, Z., Zehua, L., Tong, S., and Zhi, C. (2021). Ncapg, mediated by mir-378a-3p, regulates cell proliferation, cell cycle progression, and apoptosis of oral squamous cell carcinoma through the gsk-3β /β -catenin signaling. Neoplasma, 68(6).

Jiao, X., Ye, J., Wang, X., Yin, X., Zhang, G., and Cheng, X. (2019). Kiaa1199, a target of micorna-486-5p, promotes papillary thyroid cancer invasion by influencing epithelial-mesenchymal transition (emt). Medical Science Monitor: International Medical Journal of Experimental and Clinical Research, 25:6788.

Jung, H. M., Phillips, B. L., Patel, R. S., Cohen, D. M., Jakymiw, A., Kong, W. W., Cheng, J. Q., and Chan, E. K. (2012). Keratinization-associated mir-7 and mir-21 regulate tumor suppressor reversion-inducing cysteine-rich protein with kazal motifs (reck) in oral cancer. Journal of Biological Chemistry, 287(35):29261–29272.

Kang, Y., Zhang, Y., Zhang, Y., and Sun, Y. (2021). Nrp2, a potential biomarker for oral squamous cell carcinoma. American Journal of Translational Research, 13(8):8938.

Karmakar, B., Das, S., Bhattacharya, S., Sarkar, R., and Mukhopadhyay, I. (2019). Tight clustering for large datasets with an application to gene expression data. Scientific reports, 9(1):3053.

Khan, P., Siddiqui, J. A., Kshirsagar, P. G., Venkata, R. C., Maurya, S. K., Mirzapoiazova, T., Perumal, N., Chaudhary, S., Kanchan, R. K., Fatima, M., et al. (2023). Microrna-1 attenuates the growth and metastasis of small cell lung cancer through cxcr4/foxm1/rrm2 axis. Molecular cancer, 22(1):1.

Koner, S., Sarkar, N. D., and Laha, N. (2023). mirna-biomarker. https://github.com/SalilKoner/miRNA-biomarker.

Korthauer, K., Kimes, P. K., Duvallet, C., Reyes, A., Subramanian, A., Teng, M., Shukla, C., Alm, E. J., and Hicks, S. C. (2019). A practical guide to methods controlling false discoveries in computational biology. Genome biology, 20(1):1–21.

Koutná, I., Krontorád, P., Svoboda, Z., Bártová, E., Kozubek, M., and Kozubek, S. (2007). New insights into gene positional clustering and its properties supported by large-scale analysis of various differentiation pathways. Genomics, 89(1):81–88.

Lähnemann, D., Köster, J., Szczurek, E., McCarthy, D. J., Hicks, S. C., Robinson, M. D., Vallejos, C. A., Campbell, K. R., Beerenwinkel, N., Mahfouz, A., et al. (2020). Eleven grand challenges in single-cell data science. Genome biology, 21(1):1–35.

Leone, V., D’Angelo, D., Rubio, I., de Freitas, P. M., Federico, A., Colamaio, M., Pallante, P., Medeiros-Neto, G., and Fusco, A. (2011). MiR-1 Is a Tumor Suppressor in Thyroid Carcinogenesis Targeting CCND2, CXCR4, and SDF-1Œ±. The Journal of Clinical Endocrinology & Metabolism, 96(9):E1388– E1398.

Li, A. and Barber, R. F. (2019). Multiple testing with the structure-adaptive benjamini–hochberg algorithm. Journal of the Royal Statistical Society: Series B (Statistical Methodology), 81(1):45–74.

Li, Q., Sun, Q., and Zhu, B. (2020). Lncrna xist inhibits the progression of oral squamous cell carcinoma via sponging mir-455-3p/btg2 axis. OncoTargets and therapy, pages 11211–11220.

Li, Z., Meng, Q., Pan, A., Wu, X., Cui, J., Wang, Y., and Li, L. (2016). Microrna-455-3p promotes invasion and migration in triple negative breast cancer by targeting tumor suppressor ei24. Oncotarget, 8(12):19455.

Lin, X., Wu, W., Ying, Y., Luo, J., Xu, X., Zheng, L., Wu, W., Yang, S., and Zhao, S. (2022). Microrna-31: a pivotal oncogenic factor in oral squamous cell carcinoma. Cell Death Discovery, 8(1):140.

Liu, A., Liu, L., and Lu, H. (2019). Lncrna xist facilitates proliferation and epithelial–mesenchymal transition of colorectal cancer cells through targeting mir-486-5p and promoting neuropilin-2. Journal of cellular physiology, 234(8):13747–13761.

Liu, A., Zhu, J., Wu, G., Cao, L., Tan, Z., Zhang, S., Jiang, L., Wu, J., Li, M., Song, L., et al. (2017). Antagonizing mir-455-3p inhibits chemoresistance and aggressiveness in esophageal squamous cell carcinoma. Molecular cancer, 16(1):106.

Liu, H., Sun, M., Du, D., Pan, H., Cheng, T., Wang, J., and Zhang, Q. (2015). Whole-transcriptome analysis of differentially expressed genes in the vegetative buds, floral buds and buds of chrysanthemum morifolium. PLoS One, 10(5):e0128009.

Liu, K. Y. P., Zhu, S. Y., Brooks, D., Bowlby, R., Durham, J. S., Ma, Y., Moore, R. A., Mungall, A. J., Jones, S., and Poh, C. F. (2020). Tumor microrna profile and prognostic value for lymph node metastasis in oral squamous cell carcinoma patients. Oncotarget, 11(23):2204.

Liu, L., Guo, B., Han, Y., Xu, S., and Liu, S. (2022a). March1 silencing suppresses growth of oral squamous cell carcinoma through regulation of phlpp2. Clinical and Translational Oncology, 24(7):1311– 1321.

Liu, X., Ma, X., Li, H., Wang, Y., Mao, M., Liang, C., and Hu, Y. (2022b). Linc00472 suppresses oral squamous cell carcinoma growth by targeting mir-455-3p/elf3 axis. Bioengineered, 13(1):1162–1173.

Lu, Z., He, Q., Liang, J., Li, W., Su, Q., Chen, Z., Wan, Q., Zhou, X., Cao, L., Sun, J., et al. (2019). mir-31-5p is a potential circulating biomarker and therapeutic target for oral cancer. Molecular Therapy-Nucleic Acids, 16:471–480.

Luecken, M. D. and Theis, F. J. (2019). Current best practices in single-cell rna-seq analysis: a tutorial. Molecular systems biology, 15(6):e8746.

Manikandan, M., Deva Magendhra Rao, A.K., Arunkumar, G., Manickavasagam, M., Rajkumar, K. S., Rajaraman, R., and Munirajan, A. K. (2016). Oral squamous cell carcinoma: microrna expression profiling and integrative analyses for elucidation of tumourigenesis mechanism. Molecular cancer, 15(1):28.

Meng, W., Ye, Z., Cui, R., Perry, J., Dedousi-Huebner, V., Huebner, A., Wang, Y., Li, B., Volinia, S., Nakanishi, H., et al. (2013). Microrna-31 predicts the presence of lymph node metastases and survival in patients with lung adenocarcinomamirna-31 in lung adenocarcinoma. Clinical cancer research, 19(19):5423–5433.

Menyhart, O., Weltz, B., and Győrffy, B. (2021). Multipletesting.com: A tool for life science researchers for multiple hypothesis testing correction. PLOS ONE, 16(1):e0245824.

Min, A., Zhu, C., Peng, S., Rajthala, S., Costea, D. E., and Sapkota, D. (2015). Micrornas as important players and biomarkers in oral carcinogenesis. BioMed research international, 2015(1):186904.

Nakagawa, S., Johnson, P. C., and Schielzeth, H. (2017). The coefficient of determination r2 and intra-class correlation coefficient from generalized linear mixed-effects models revisited and expanded. Journal of the Royal Society Interface, 14(134):20170213.

Nohata, N., Hanazawa, T., Kikkawa, N., Sakurai, D., Sasaki, K., Chiyomaru, T., Kawakami, K., Yoshino, H., Enokida, H., Nakagawa, M., et al. (2011a). Identification of novel molecular targets regulated by tumor suppressive mir-1/mir-133a in maxillary sinus squamous cell carcinoma. International journal of oncology, 39(5):1099–1107.

Nohata, N., Sone, Y., Hanazawa, T., Fuse, M., Kikkawa, N., Yoshino, H., Chiyomaru, T., Kawakami, K., Enokida, H., Nakagawa, M., et al. (2011b). mir-1 as a tumor suppressive microrna targeting tagln2 in head and neck squamous cell carcinoma. Oncotarget, 2(1-2):29.

Okada, R., Koshizuka, K., Yamada, Y., Moriya, S., Kikkawa, N., Kinoshita, T., Hanazawa, T., and Seki, N. (2019). Regulation of oncogenic targets by miR-99a-3p (passenger strand of mir-99a-duplex) in head and neck squamous cell carcinoma. Cells, 8(12):1535.

Olasz, E. B., Seline, L. N., Schock, A. M., Duncan, N. E., Lopez, A., Lazar, J., Flister, M. J., Lu, Y., Liu, P., Sokumbi, O., et al. (2015). Microrna-135b regulates leucine zipper tumor suppressor 1 in cutaneous squamous cell carcinoma. PloS one, 10(5):e0125412.

Osako, Y., Yoshino, H., Sakaguchi, T., Sugita, S., Yonemori, M., Nakagawa, M., and Enokida, H. (2019). Potential tumor-suppressive role of microrna-99a-3p in sunitinib-resistant renal cell carcinoma cells through the regulation of rrm2. International journal of oncology, 54(5):1759–1770.

Osan, C., Chira, S., Nutu, A. M., Braicu, C., Baciut, M., Korban, S. S., and Berindan-Neagoe, I. (2021). The connection between micrornas and oral cancer pathogenesis: emerging biomarkers in oral cancer management. Genes, 12(12):1989.

Pansare, K., Gardi, N., Kamat, S., Dange, P., Previn, R., Gera, P., Kowtal, P., Amin, K., and Sarin, R. (2019). Establishment and genomic characterization of gingivobuccal carcinoma cell lines with smokeless tobacco associated genetic alterations and oncogenic pik3ca mutation. Scientific reports, 9(1):1–10.

Peng, C.-Y., Liao, Y.-W., Lu, M.-Y., Yu, C.-H., Yu, C.-C., and Chou, M.-Y. (2017). Downregulation of miR-1 enhances tumorigenicity and invasiveness in oral squamous cell carcinomas. Journal of the Formosan Medical Association, 116(10):782–789.

Rajan, C., Roshan, V. D., Khan, I., Manasa, V., Himal, I., Kattoor, J., Thomas, S., Kondaiah, P., and Kannan, S. (2021). Mirna expression profiling and emergence of new prognostic signature for oral squamous cell carcinoma. Scientific reports, 11(1):7298.

Reddy, C. S. S., Pp, A. S. U., Ganapathy, D. M., Kp, A., and Sekar, D. (2024). Microrna-21 as a biomarker in terminal stage oral squamous cell carcinoma (oscc) in the south indian population. Oral Oncology Reports, 9:100139.

Reis, P. P., Tomenson, M., Cervigne, N. K., Machado, J., Jurisica, I., Pintilie, M., Sukhai, M. A., Perez-Ordonez, B., Grénman, R., Gilbert, R. W., et al. (2010). Programmed cell death 4 loss increases tumor cell invasion and is regulated by miR-21 in oral squamous cell carcinoma. Molecular cancer, 9(1):238.

Safa, A., Bahroudi, Z., Shoorei, H., Majidpoor, J., Abak, A., Taheri, M., and Ghafouri-Fard, S. (2020). mir-1: A comprehensive review of its role in normal development and diverse disorders. Biomedicine & Pharmacotherapy, 132:110903.

Sarkar, S. K. and Zhao, Z. (2022). Local false discovery rate based methods for multiple testing of one-way classified hypotheses. Electronic Journal of Statistics, 16(2):6043–6085.

Schneider, A., Victoria, B., Lopez, Y. N., Suchorska, W., Barczak, W., Sobecka, A., Golusinski, W., Masternak, M. M., and Golusinski, P. (2018). Tissue and serum microrna profile of oral squamous cell carcinoma patients. Scientific reports, 8(1):675.

Searle, S. R., Casella, G., and McCulloch, C. E. (2009). Variance components. John Wiley & Sons.

Sesia, M., Bates, S., Candés, E., Marchini, J., and Sabatti, C. (2021a). False discovery rate control in genome-wide association studies with population structure. Proceedings of the National Academy of Sciences, 118(40):e2105841118.

Sesia, M., Bates, S., Candés, E., Marchini, J., and Sabatti, C. (2021b). False discovery rate control in genome-wide association studies with population structure. Proceedings of the National Academy of Sciences, 118(40):e2105841118.

Shao, Y., Zhang, S., Pan, Y., Peng, Z., and Dong, Y. (2025). mir-135b: A key role in cancer biology and therapeutic targets. Non-coding RNA Research, 12:67.

Singh, R., De Sarkar, N., Sarkar, S., Roy, R., Chattopadhyay, E., Ray, A., Biswas, N. K., Maitra, A., and Roy, B. (2017). Analysis of the whole transcriptome from gingivo-buccal squamous cell carcinoma reveals deregulated immune landscape and suggests targets for immunotherapy. PloS one, 12(9):e0183606.

Soga, D., Yoshiba, S., Shiogama, S., Miyazaki, H., Kondo, S., and Shintani, S. (2013). microrna expression profiles in oral squamous cell carcinoma. Oncology reports, 30(2):579–583.

Storey, J. D. (2002). A direct approach to false discovery rates. Journal of the Royal Statistical Society Series B: Statistical Methodology, 64(3):479–498.

Troiano, G., Mastrangelo, F., Caponio, V., Laino, L., Cirillo, N., and Lo Muzio, L. (2018). Predictive prognostic value of tissue-based microrna expression in oral squamous cell carcinoma: a systematic review and meta-analysis. Journal of dental research, 97(7):759–766.

Wang, J., Wang, Y., Sun, D., Bu, J., Ren, F., Liu, B., Zhang, S., Xu, Z., Pang, S., and Xu, S. (2017). miR-455-5p promotes cell growth and invasion by targeting soco3 in non-small cell lung cancer. Oncotarget, 8(70):114956.

Wang, Y., Huang, D., Li, M., and Yang, M. (2025). Microrna-99 family in cancer: molecular mechanisms for clinical applications. PeerJ, 13:e19188.

Wang, Y., Shang, S., Yu, K., Sun, H., Ma, W., and Zhao, W. (2020). miR-224, miR-147b and miR-31 associated with lymph node metastasis and prognosis for lung adenocarcinoma by regulating PRPF4B, WDR82 or NR3C2. PeerJ, 8:e9704.

Wei, G.-G., Guo, W.-P., Tang, Z.-Y., Li, S.-H., Wu, H.-Y., and Zhang, L.-C. (2019a). Expression level and prospective mechanism of mirna-99a-3p in head and neck squamous cell carcinoma based on mirna-chip and mirna-sequencing data in 1, 167 cases. Pathology-Research and Practice, 215(5):963–976.

Wei, Q., Yao, J., and Yang, Y. (2019b). Microrna-1247 inhibits the viability and metastasis of osteosarcoma cells via targeting nrp1 and mediating wnt/β -catenin pathway. Eur Rev Med Pharmacol Sci, 23(17):7266–7274.

Wong, N., Khwaja, S. S., Baker, C. M., Gay, H. A., Thorstad, W. L., Daly, M. D., Lewis Jr, J. S., and Wang, X. (2016). Prognostic micro rna signatures derived from the cancer genome atlas for head and neck squamous cell carcinomas. Cancer medicine, 5(7):1619–1628.

Wu, J., Li, Y., Liu, J., and Xu, Y. (2021). Down-regulation of lncrna hcg11 promotes cell proliferation of oral squamous cell carcinoma through sponging mir-455-5p. The Journal of Gene Medicine, 23(3):e3293.

Yamamoto, N., Nishikawa, R., Chiyomaru, T., Goto, Y., Fukumoto, I., Usui, H., Mitsuhashi, A., Enokida, H., Nakagawa, M., Shozu, M., et al. (2015). The tumor-suppressive microrna-1/133a cluster targets pde7a and inhibits cancer cell migration and invasion in endometrial cancer. International journal of oncology, 47(1):325–334.

Yan, Y., Wang, X., Venø, M. T., Bakholdt, V., Sørensen, J. A., Krogdahl, A., Sun, Z., Gao, S., and Kjems, J. (2016). Circulating mirnas as biomarkers for oral squamous cell carcinoma recurrence in operated patients. Oncotarget, 8(5):8206.

Yang, H., He, C., Feng, Y., and Jin, J. (2023). Exosome-delivered mir-486-3p inhibits the progression of osteosarcoma via sponging circkeap1/march1 axis components. Oncology Letters, 27(1):24.

Yang, H., Huang, Y., He, J., Chai, G., Di, Y., Wang, A., and Gui, D. (2020). Mir-486-3p inhibits the proliferation, migration and invasion of retinoblastoma cells by targeting ecm1. Bioscience Reports, 40(6).

Ye, L., Fan, T., Qin, Y., Qiu, C., Li, L., Dai, M., Zhou, Y., Chen, Y., and Jiang, Y. (2023). Microrna-455-3p accelerate malignant progression of tumor by targeting h2afz in colorectal cancer. Cell Cycle, 22(7):777–795.

Yi, J. M., Kang, E.-J., Kwon, H.-M., Bae, J.-H., Kang, K., Ahuja, N., and Yang, K. (2017). Epigenetically altered mir-1247 functions as a tumor suppressor in pancreatic cancer. Oncotarget, 8(16):26600.

Zhang, B., Zhou, Y., Lin, N., Lowdon, R. F., Hong, C., Nagarajan, R. P., Cheng, J. B., Li, D., Stevens, M., Lee, H. J., et al. (2013). Functional dna methylation differences between tissues, cell types, and across individuals discovered using the m&m algorithm. Genome research, 23:1522–1540.

Zhang, J., Fu, J., Pan, Y., Zhang, X., and Shen, L. (2016a). Silencing of mir-1247 by dna methylation promoted non-small-cell lung cancer cell invasion and migration by effects of stmn1. OncoTargets and therapy, 9:7297.

Zhang, J., Fu, J., Pan, Y., Zhang, X., and Shen, L. (2016b). Silencing of mir-1247 by dna methylation promoted non-small-cell lung cancer cell invasion and migration by effects of stmn1. OncoTargets and therapy, pages 7297–7307.

Zhang, W. C., Wells, J. M., Chow, K.-H., Huang, H., Yuan, M., Saxena, T., Melnick, M. A., Politi, K., Asara, J. M., Costa, D. B., et al. (2019). miR-147b-mediated TCA cycle dysfunction and pseudohypoxia initiate drug tolerance to egfr inhibitors in lung adenocarcinoma. Nature metabolism, 1(4):460–474.

Zhang, Z. (2016). Missing data imputation: focusing on single imputation. Annals of translational medicine, 4(1).

