## Supplement for "False Discovery Rate Control for Grouped Hypotheses: Application to miRNAome Data"

#### ABSTRACT

This document contains supplementary material and does not include an abstract.

#### S1 MIRNAOME DATA GENERATION

The miRNA profiling was performed using the TaqMan Low-Density Array (TLDA), a high-throughput qPCR assay suited for quantifying standard and low-expressing miRNAs in biospecimens (Wang et al., 2011). RNA was isolated using the mirVana kit (Life Technologies, USA), and only samples with RNA integrity number (RIN) $\geq 6.9$ were included. Ct, $\Delta$Ct, and $\Delta\Delta$Ct values were computed using the SDS and DataAssist software packages (Life Technologies, USA). The definitions are as follows. Ct is the cycle number at which PCR product reaches a defined threshold. $\Delta$Ct is the difference between the Ct of a miRNA and the geometric mean Ct of the three most stable endogenous control miRNAs in that tissue. $\Delta\Delta$Ct is the difference between a miRNA’s $\Delta$Ct in tumor and in matched control tissue. The raw data were normalized and trimmed following the standard TLDA data analysis protocol. Further details on the data collection methods are available in (De Sarkar et al., 2014).

#### S2 MISSINGNESS

Figure S2.1 shows that the number of missing miRNA pairs can range from 12 to 232 across the eighteen patients. Only two patients exhibit more than 100 missing pairs, while the majority, 70% of the patients, have fewer than 50 missing pairs. This histogram in Figure S2.2 indicates that a significant proportion, approximately 59.77%, of the miRNAs possess at least one pair of missing observations. However, most of these miRNAs (about 48.76% of them) have at most two missing pairs of observations. Nevertheless, there are fifteen miRNAs with 50% missing observations each.

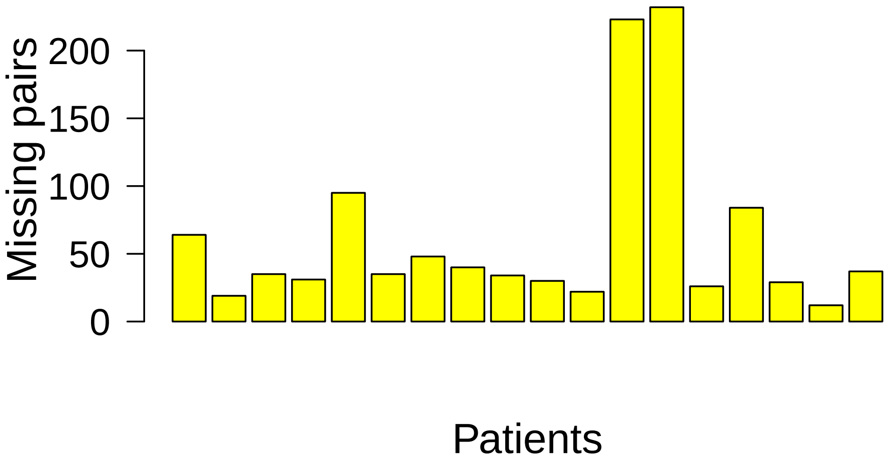


**Figure S2.1. Barplot of the number of missing pairs of miRNAs across patients**. Here the number of patients is 18, and the total number of miRNAs is 522. The x-axis corresponds to the patients, and the y-axis corresponds to the number of missing pairs of miRNAs corresponding to each patient. Each patient had at least twelve missing pairs of miRNA expressions.

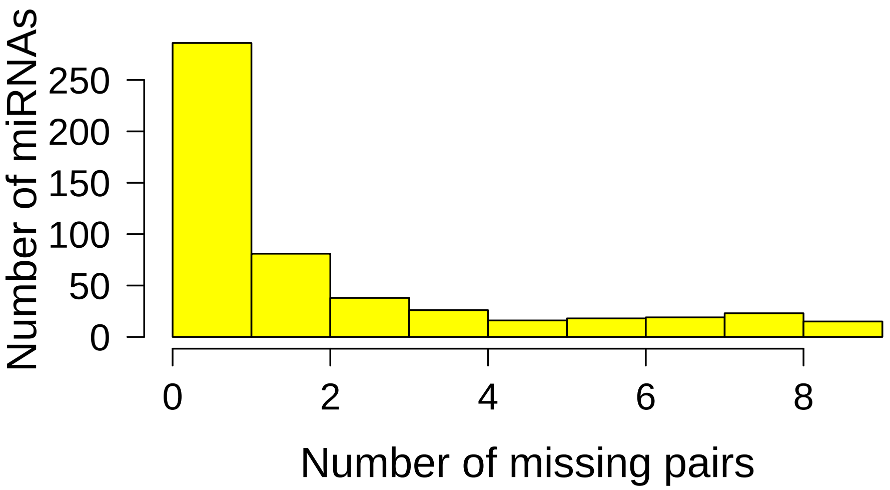


**Figure S2.2. Histogram of the number of missing pairs of miRNAs**. Here we considered the 522 miRNAs from our analysis. The x-axis corresponds to the number of missing pairs, and the y-axis corresponds to the number of miRNAs with that many missing pairs.

#### S3 PLOTS AND TABLES RELATED TO THE GROUPS

**Table S3.1. Table of group composition for Scheme (c).** This table lists the size, chromosome number, strand, and arm for each group in Scheme (c). Here size refers to the number of miRNAs in a group. The groups are ordered according to their Chromosome number. Some groups are situated on both p and q arms because single-miRNA groups were fused with the adjacent group to make a larger group.

| Group number | Size | Chromosome number | Strand | Arm |
|---|---|---|---|---|
| 1 | 9 | 1 | - | p |
| 2 | 19 | 1 | - | q |
| 3 | 10 | 1 | + | p |
| 4 | 3 | 1 | + | q |
| 5 | 3 | 2 | - | p |
| 6 | 3 | 2 | - | q |
| 7 | 8 | 2 | + | q |
| 8 | 8 | 3 | - | p |
| 9 | 2 | 3 | - | q |
| 10 | 4 | 3 | + | p |
| 11 | 10 | 3 | + | q |
| 12 | 3 | 4 | - | p |
| 13 | 6 | 4 | - | q |
| 14 | 4 | 4 | + | p |
| 15 | 5 | 4 | + | q |
| 16 | 2 | 5 | - | p |
| 17 | 9 | 5 | - | q |
| 18 | 7 | 5 | + | q |
| 19 | 4 | 6 | - | p, q |
| 20 | 2 | 6 | + | p |
| 21 | 6 | 7 | - | p |
| 22 | 17 | 7 | - | q |
| 23 | 2 | 7 | + | p |
| 24 | 7 | 7 | + | q |
| 25 | 5 | 8 | - | p |
| 26 | 8 | 8 | - | q |
| 27 | 3 | 8 | + | p, q |
| 28 | 3 | 9 | - | p |
| 29 | 4 | 9 | - | q |
| 30 | 14 | 9 | + | p, q |
| 31 | 2 | 10 | - | p |
| 32 | 4 | 10 | - | q |
| 33 | 2 | 10 | + | p |
| 34 | 4 | 10 | + | q |
| 35 | 4 | 11 | - | p |
| 36 | 11 | 11 | - | q |
| 37 | 6 | 11 | + | p, q |
| 38 | 8 | 12 | - | p, q |
| 39 | 5 | 12 | + | p |
| 40 | 8 | 12 | + | q |
| 41 | 5 | 13 | - | q |
| 42 | 13 | 13 | + | q |
| 43 | 4 | 14 | - | q |
| 44 | 55 | 14 | + | q |

Continued on next page

**Table S3.1 – continued from previous page**

| Group number | Size | Chromosome number | Strand | Arm |
|---|---|---|---|---|
| 45 | 9 | 15 | - | q |
| 46 | 7 | 15 | + | q |
| 47 | 2 | 16 | - | p, q |
| 48 | 5 | 16 | + | p |
| 49 | 3 | 16 | + | q |
| 50 | 13 | 17 | - | p |
| 51 | 17 | 17 | - | q |
| 52 | 2 | 17 | + | p |
| 53 | 5 | 17 | + | q |
| 54 | 3 | 18 | - | q |
| 55 | 8 | 19 | - | p |
| 56 | 3 | 19 | - | q |
| 57 | 5 | 19 | + | p |
| 58 | 37 | 19 | + | q |
| 59 | 3 | 20 | - | p, q |
| 60 | 7 | 20 | + | p |
| 61 | 6 | 21 | + | q |
| 62 | 4 | 22 | - | q |
| 63 | 10 | 22 | + | q |
| 64 | 6 | X | - | p |
| 65 | 32 | X | - | q |
| 66 | 9 | X | + | p |
| 67 | 5 | X | + | q |

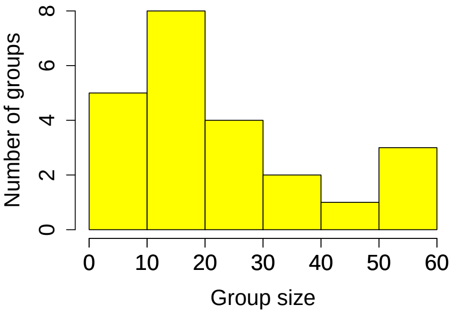


**(i) Grouped by chromosomes only (Scheme (a)).** Max size: 59, min: 3.

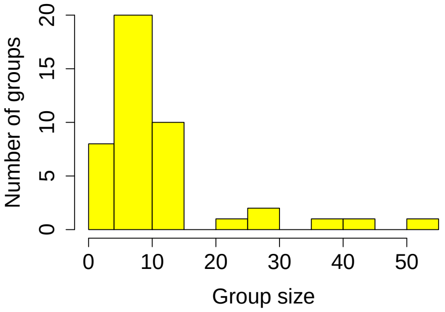


**(ii) Grouped by chromosome + strand (Scheme (b)).** Max size: 55, min: 2.

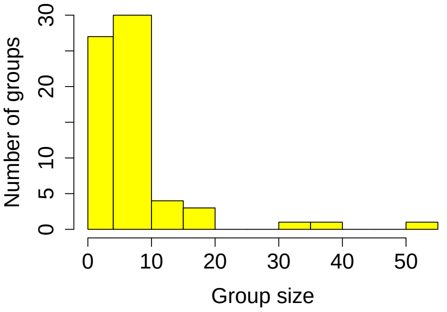


**(iii) Grouped by chromosome + strand + arm (Scheme (c)).** Max size: 55, min: 2.

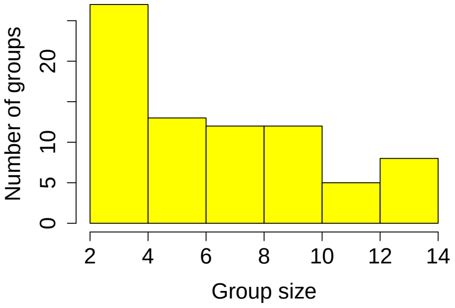


**(iv) Group size when** $k = 15$. Max size: 15, min: 2.

S3.1iv

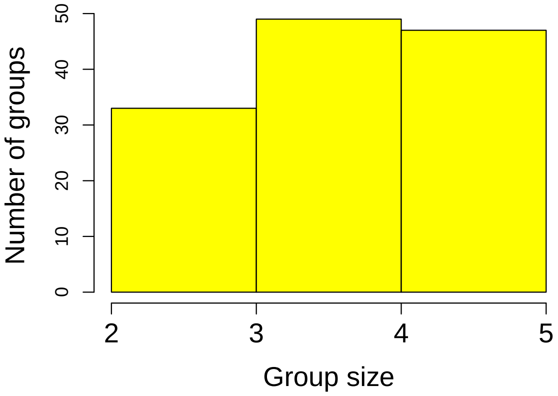


**(v) Group size when** $k = 5$. Max size: 5, min: 2.

**Figure S3.1. Histograms of group sizes under different grouping schemes.** Group size refers to the number of miRNAs in a group. Panels (i)–(iii) correspond to grouping Schemes (a), (b), and (c), respectively, which represent grouping by: (a) chromosome number only, (b) chromosome number and strand, and (c) chromosome number, strand, and arm. In panel (iv), groups from scheme (c) were further split into adjacent subgroups of at most 15 miRNAs using the miRNAs' chromosome coordinates from De Sarkar et al. (2014); panel (v) applies the same procedure with a maximum group size of 5. The x-axis shows group size, and the y-axis shows the number of groups. For each scheme, "max" and "min" indicate the largest and smallest group sizes.

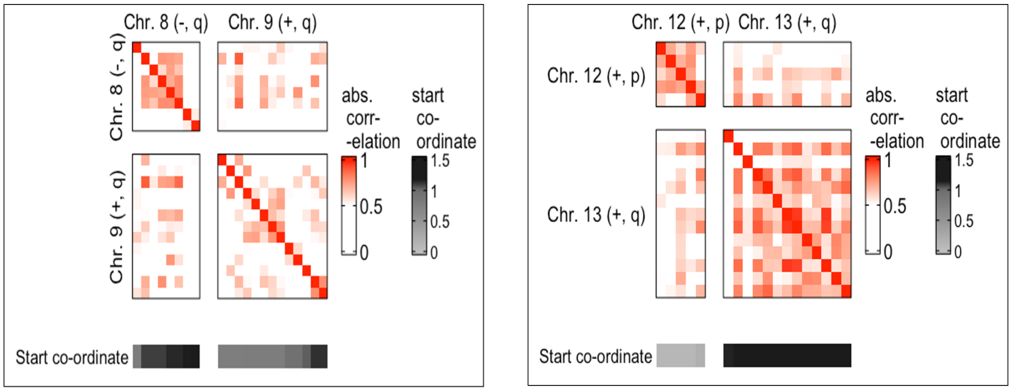


**Figure S3.2. Heatmap of the absolute correlations between the ΔΔCt values of the miRNAs from two groups.** In the left panel, the first group contains miRNAs from the q arm of the positive strand of chromosome 9 and the second group contains miRNAs from the q arm of the negative strand of chromosome 8. In the right panel, the first group corresponds to the p arm of the positive strand of chromosome 12 and the second group corresponds to the miRNAs on the q arm of the positive strand of chromosome 13. The diagonal blocks provide the heatmaps of the absolute correlation between the ΔΔCt values of miRNAs within the same group, whereas the off-diagonal blocks showcase the absolute correlation between ΔΔCt values of miRNAs from different groups. Only absolute correlations above 0.5 are presented in the heatmaps. The miRNAs are ordered based on their start coordinates on the chromosome, ensuring that adjacent miRNAs are positioned next to each other in the heatmap. The sidebars below the heatmaps provide the start coordinates of the miRNAs.

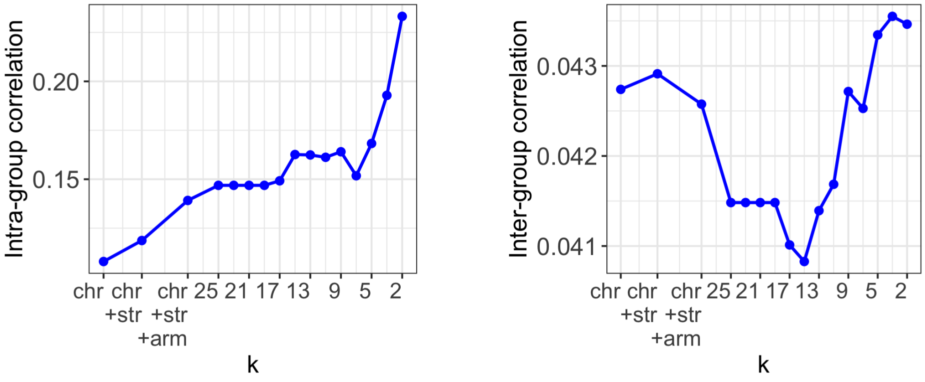


**(i)** Inter-group correlation vs k **(ii)** Inter-group correlations vs k

**Figure S3.3. Plot of Intra- and Inter-group correlations.** The x-axis corresponds to different grouping schemes, and the Y-axis represents the correlation estimates for each scheme. For each grouping scheme, the intra- and inter-group correlations are computed according to the simple random effect model discussed in Appendix 5. The grouping schemes were as follows: Chr refers to Scheme (a) or grouping by chromosome number only; Chr+Str refers to Scheme (b) or grouping by chromosome number and strand; and Chr+Str+Arm refers to Scheme (c) or grouping by chromosome number, strand, and arm. For each $k$, any group from Scheme (c) with more than $k$ miRNAs was split into adjacent groups of size $k$. If the group size was not a multiple of $k$, the final group had fewer than $k$ miRNAs. The miRNAs' coordinates on the chromosome, provided by De Sarkar et al. (2014), were used to guide this splitting so that adjacent miRNAs were grouped together. Here, apart from the $k$'s used in the main manuscript, i.e., $k \in \{25, 23, \ldots, 5\}$, we also consider $k = 2$ and 3 to highlight how the intra-group correlation increases in smaller groups.

### S4 TABLES AND FIGURES CORRESPONDING TO SECTION 3

| miRNA | Chr. | Strand | Arm | ΔΔCt | p-value | Missing pairs | BH | TST | LSL | SABHA |
|---|---|---|---|---|---|---|---|---|---|---|
| hsa-miR-133a-3p* | 18 | - | q | 6.7 | $5.32E^{-05}\downarrow$ | 0 | x | x | x | x |
| hsa-miR-31-3p* | 9 | - | p | -3.8 | $1.31E^{-04}\uparrow$ | 2 | x | x | x | x |
| hsa-miR-206* | 6 | + | p | 6.0 | $1.97E^{-04}\downarrow$ | 0 | x | x | x | x |
| hsa-miR-31-5p* | 9 | - | q | -3.4 | $6.18E^{-04}\uparrow$ | 0 | | x | (a) | |
| hsa-miR-204-5p* | 9 | - | q | 4.6 | $8.81E^{-04}\downarrow$ | 2 | | x | x | |
| hsa-miR-1 | 18 | - | q | 5.2 | $9.38E^{-04}\downarrow$ | 0 | | x | x | x |
| hsa-miR-7-5p* | 15 | + | q | -3.1 | $9.50E^{-04}\uparrow$ | 2 | | x | x | |
| hsa-miR-1293* | 12 | - | q | -4.8 | $2.77E^{-03}\uparrow$ | 9 | | (b) | | |
| hsa-miR-486-3p | 8 | - | p | 2.4 | $2.81E^{-03}\downarrow$ | 0 | | x | | |
| hsa-miR-21-5p | 17 | + | q | -2.2 | $3.67E^{-03}\uparrow$ | 0 | | (b) | | |
| hsa-miR-147b | 15 | + | q | -2.1 | $5.52E^{-03}\uparrow$ | 2 | | x | | |
| hsa-miR-99a-3p | 21 | + | q | 2.8 | $5.83E^{-03}\downarrow$ | 0 | | x | | |
| hsa-miR-455-5p | 9 | + | q | -1.4 | $5.98E^{-03}\uparrow$ | 0 | | (a) | | |
| hsa-miR-1247-5p | 14 | - | q | 2.4 | $8.93E^{-03}\downarrow$ | 2 | | (b) | | |

**Table S4.1. Significantly deregulated miRNAs for grouping Schemes (a) and (b).** The listed miRNAs were detected as significantly deregulated (5% significance level) by BH, TST-GBH (TST), LSL-GBH (LSL), or SABHA under either Scheme (a) or Scheme (b). In Scheme (a), the miRNAs were grouped based on shared chromosome number. In Scheme (b), the groupings are based on shared chromosome number and strand. If a method detected a miRNA with both grouping schemes, the corresponding row is marked by x. Otherwise, the row marks the grouping scheme under which the corresponding miRNA were detected. The "Chr." column indicates the chromosome number; the "strand" column indicates the transcribing strand; and the "arm" column indicates the arm of the miRNA gene. The "ΔΔCt" column reports the average ΔΔCt value across 18 patients for each miRNA, serving as a surrogate estimate of differential expression. As TST-GBH, LSL-GBH, and SABHA do not yield FDR-corrected p-values, we have reported the raw p-values before FDR correction. The rows are sorted according to p-value. The arrow signs after the p-values indicate whether the expression of the miRNAs was upregulated (↑) or downregulated (↓) (per the sign of ΔΔCt). The "Missing pairs" column shows the number of missing observations for each miRNA. The expressions of miRNAs superscripted with an asterisk were reported to be deregulated by De Sarkar et al. (2014).

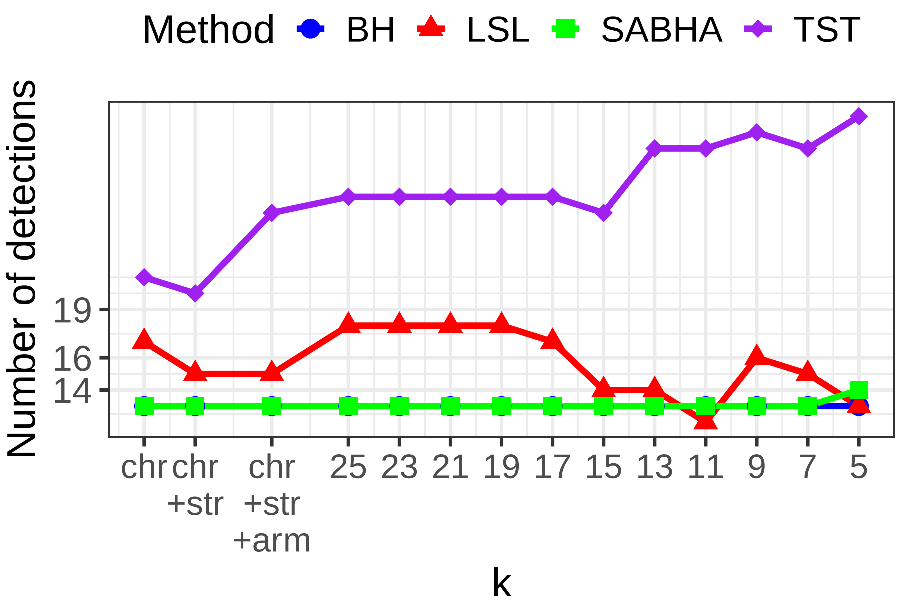


**Figure S4.1. Plot of the number of detections by each FDR control method vs grouping scheme for the median-imputed data.** The methods compared were BH, TST-GBH (abbreviated as TST), LSL-GBH (abbreviated as LSL), and SABHA. The first three grouping schemes were: (a) grouping by chromosome number only (Chr); (b) grouping by chromosome number and strand (Chr+strand); and (c) grouping by chromosome number, strand, and arm (Chr+Str+Arm). For each *k*, any group from Scheme (c) with more than *k* miRNAs was split into adjacent groups of size *k*, using the miRNAs' chromosome coordinates from De Sarkar et al. (2014) to ensure adjacent miRNAs were grouped together. If the group size was not a multiple of *k*, the final group contained fewer than *k* miRNAs.The Y-axis shows the total number of detections from each grouping scheme.

| MiRNA | chr | strand | Arm | ΔΔ*Ct* | p-value | Missing pairs | BH | TST | LSL | SABHA |
|---|---|---|---|---|---|---|---|---|---|---|
| hsa-miR-1293 | 12 | - | q | -4.8 | 1.51E-06 ↑ | 9 | x | x | | x |
| hsa-miR-31-3p | 9 | - | p | -3.8 | 2.00E-05 ↑ | 2 | x | x | x | x |
| hsa-miR-548k† | 11 | + | p | -2.7 | 3.86E-05 ↑ | 8 | x | x | | x |
| hsa-miR-133a-3p | 18 | - | q | 6.7 | 5.32E-05 ↓ | 0 | x | x | x | x |
| hsa-miR-206 | 6 | + | p | 6.0 | 1.97E-04 ↓ | 0 | x | x | | x |
| hsa-miR-204-5p | 9 | - | q | 4.6 | 2.19E-04 ↓ | 2 | x | x | | x |
| hsa-miR-7-5p | 15 | + | q | -3.1 | 2.80E-04 ↑ | 2 | x | x | x | x |
| hsa-miR-891a-5p† | X | - | q | 4.4 | 2.84E-04 ↓ | 7 | x | x | x | x |
| hsa-miR-504-5p† | X | - | q | 4.7 | 6.00E-04 ↓ | 6 | x | x | x | x |
| hsa-miR-31-5p | 9 | - | p | 3.4 | 6.18E-04 ↑ | 0 | x | x | x | x |
| hsa-miR-133b† | 6 | + | p | 2.7 | 6.80E-04 ↓ | 8 | x | x | | x |
| hsa-miR-508-3p† | X | - | q | 4.0 | 7.91E-04 ↓ | 6 | x | x | x | x |
| hsa-miR-1 | 18 | - | q | 5.2 | 9.38E-04 ↓ | 0 | x | x | x | x |
| hsa-miR-135b-3p† | 1 | - | q | -2.2 | 2.42E-03 ↑ | 2 | | x | | |
| hsa-miR-200b-5p† | 1 | + | p | -3.5 | 2.66E-03 ↑ | 3 | | x | | |
| hsa-miR-486-3p | 8 | - | p | 2.4 | 2.81E-03 ↓ | 0 | | x | x | |
| hsa-miR-147b | 15 | + | q | -2.1 | 2.96E-03 ↑ | 2 | | x | x | |
| hsa-miR-1290† | 1 | - | p | -2.2 | 3.45E-03 ↑ | 2 | | x | | |
| hsa-miR-299-5p† | 14 | + | q | 2.5 | 3.62E-03 ↓ | 4 | | | x | |
| hsa-miR-21-5p | 17 | + | q | -2.2 | 3.66E-03 ↑ | 0 | | x | | |
| hsa-miR-770-5p† | 14 | + | q | 3.0 | 3.71E-03 ↓ | 3 | | | x | |
| hsa-miR-1247-5p | 14 | - | q | 2.4 | 4.40E-03 ↓ | 2 | | x | x | |
| hsa-miR-211-5p† | 15 | - | q | 3.7 | 4.90E-03 ↓ | 4 | | x | x | |
| hsa-miR-200c-5p† | 12 | + | p | -2.3 | 5.31E-03 ↑ | 3 | | x | | |
| hsa-miR-99a-3p | 21 | + | q | 2.8 | 5.83E-03 ↓ | 0 | | x | x | |
| hsa-miR-486-5p | 8 | - | p | 2.0 | 1.78E-02 ↓ | 0 | | x | | |
| hsa-miR-383-5p† | 8 | - | p | 2.7 | 3.05E-02 ↓ | 4 | | x | | |

**Table S4.2. Significantly deregulated miRNAs detected using the median-imputed data.** The listed miRNAs were detected as significantly deregulated (5% significance level) by BH, TST-GBH (TST), LSL-GBH (LSL), or SABHA under either Scheme (a) or Scheme (b). In Scheme (a), the miRNAs were grouped based on shared chromosome number. In Scheme (b), the groupings are based on shared chromosome number and strand. If a method detected a miRNA with both grouping schemes, the corresponding row is marked by x. The "Chr." column indicates the chromosome number; the "strand" column indicates the transcribing strand; and the "arm" column indicates the arm of the miRNA gene. The ΔΔCt values are used as surrogate estimates of differential expression for each miRNA. As TST-GBH, LSL-GBH, and SABHA do not yield FDR-corrected p-values, we have reported the raw p-values before FDR correction. The rows are sorted according to p-value. The arrow signs after the p-values indicate whether the expression of the miRNAs was upregulated (↑) or downregulated (↓) (per the sign of ΔΔCt). The "Missing pairs" column shows the number of missing observations for each miRNA.
† These miRNAs were not detected in the non-imputed data. In total, there are thirteen such cases, many with moderate to severe missingness, suggesting that their detection may be driven by variance deflation from imputation.

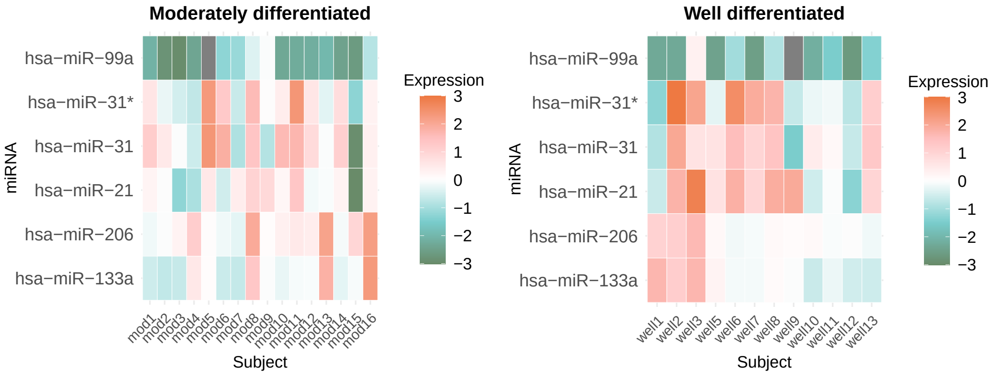


**Figure S4.2. Heatmap of Δ$Ct$ values of miRNAs in OSCC patients from Manikandan et al. (2016)'s data.** The heatmap shows the Δ$Ct$ values of miRNAs studied in Manikandan et al. (2016) that overlap with the 16 deregulated miRNAs listed in Table 1. We generated this heatmap using the dataset available from the journal website of the published study. This study compared miRNA expression in moderately differentiated and well-differentiated oral squamous cell carcinoma (OSCC) relative to normal squamous epithelial cells. Well-differentiated tumors are generally more aggressive than moderately differentiated tumors. The left and right heatmaps correspond to patients with moderately and well-differentiated tumors, respectively. Note that the Δ$Ct$ values in Manikandan et al. (2016) were normalized differently from those in De Sarkar et al. (2014), and are therefore not directly comparable. Missing observations are shown in white. hsa-miR-21-5p and hsa-miR-99a-3p are among the novel miRNAs, which were up and downregulated, respectively in De Sarkar et al. (2014)'s data. Note that miR-21's expression in the well differentiated tumors were higher compared to the moderately differentiated tumors, indicating that miR-21 may play a role in early-stage tumorigenesis.

### REFERENCES

De Sarkar, N., Roy, R., Mitra, J. K., Ghose, S., Chakraborty, A., Paul, R. R., Mukhopadhyay, I., and Roy, B. (2014). A quest for mirna bio-marker: a track back approach from gingivo buccal cancer to two different types of precancers. *PloS one*, 9(8):e104839.

Manikandan, M., Deva Magendhra Rao, A. K., Arunkumar, G., Manickavasagam, M., Rajkumar, K. S., Rajaraman, R., and Munirajan, A. K. (2016). Oral squamous cell carcinoma: microrna expression profiling and integrative analyses for elucidation of tumourigenesis mechanism. *Molecular cancer*, 15(1):28.

Wang, B., Howel, P., Bruheim, S., Ju, J., Owen, L. B., Fodstad, O., and Xi, Y. (2011). Systematic evaluation of three microrna profiling platforms: microarray, beads array, and quantitative real-time pcr array. *PloS one*, 6(2):e17167.
